# Fibroblast orchestration of inflammaging via NF-kB activation

**DOI:** 10.1101/2025.10.01.679697

**Authors:** Nancy C. Allen, Christian Ringler, Jin Young Lee, Nabora Reyes, Ritusree Biswas, Sofia Caryotakis, Melia Magnen, Pedro Ruivo, Vincent Auyeung, Mark Looney, Averil Ma, Ari B. Molofsky, Tien Peng

## Abstract

Aged tissue is characterized by chronic inflammation known as “inflammaging”. While this aging immune phenotype supposedly drives some of the most common diseases affecting the elderly, little is known about the structural drivers of inflammaging. In this study, we demonstrate that age-dependent activation of NF-kB in tissue fibroblasts remodels the immune architecture, promoting the emergence of an exhausted T cell population (GZMK^+^/CD8^+^) recently identified in normal aging, as well as autoimmunity and cancer. Fibroblast-specific NF-kB activation triggered a fibroblast-macrophage-T cell circuit to form tertiary lymphoid structures in the lung and promoted the emergence of exhausted GZMK^+^ T cells. Fibroblastic activation of NF-kB increased host susceptibility to acute lung injury and mimics severe pneumonia commonly seen in elderly patients, which was alleviated by deletion of GZMK^+^ T cells. Our data provide a structural basis for inflammaging, where fibroblasts orchestrate the complex immune aging phenotype in non-immune tissues, increasing susceptibility to age-related diseases.

**Highlights:**

- Bronchus-associated lymphoid tissue (BALT) enriched for GZMK+ T cells develop with age
- Lung adventitial fibroblasts demonstrate increased NF-kB activation with age.
- Fibroblast activation of NF-kB in young animals recapitulates multiple features of normal lung immune aging
- Depletion of GZMK+ cells decreases lung inflammation in a mouse model of acute respiratory distress syndrome (ARDS)

## Introduction

Aging drives complex changes in both the adaptive and innate immune systems that result in altered immune function in the elderly. An important part of these age-related immune changes, generally termed “immunosenescence”, is the development of a chronic proinflammatory state known as “inflamm-aging”^1,2^ . Inflammaging, initially postulated as a response to chronic antigens and cellular stress^2^ has been characterized in multiple ways, from the elevation of inflammatory biomarkers^3^ to single cell analyses that describe tissue infiltration of immune cells in aged organs^4,5^. Chronic inflammation is thought to be an important component of many common non-communicable diseases that disproportionally affect the elderly, creating an important potential link between inflammaging and age-related disease susceptibility^6^.

In addition to advanced age as a risk factor for chronic inflammatory disease, elderly individuals are also more susceptible to acute inflammatory disease, as most recently illustrated by the COVID-19 pandemic that inordinately afflicted the elderly^7^. Those who succumbed to the disease mostly died of severe pneumonia characterized by an exaggerated inflammatory response, also known as acute respiratory distress syndrome (ARDS)^8^. In contrast to the overwhelming epidemiological data that links inflammation with age-associated diseases, much less is known about the molecular and structural determinants of the inflammaging phenotype in tissues, and whether the aging-associated changes in tissue-resident immune cells might be related to the aging of structural cell types within the host organ.

Utilizing single cell/spatial transcriptomics and combinatorial mouse genetic models, we show that NF-kB-activated fibroblasts accumulating in aged tissues are critical orchestrators of the complex inflammaging phenotype. This phenotype includes the formation of tertiary lymphoid structures within the lung (known as inducible bronchus-associated lymphoid tissue, or BALT) with topographic features that induce exhausted GZMK^+^ T cells recently identified in aged tissues along with age-related pathologies^9–11^. Furthermore, we show that GZMK^+^ T cells are not merely bystanders of the aging process, but rather a potent driver of inflammaging that could be targeted to alleviate age-related inflammatory diseases. Our work provides a structural basis for how immune cells interact with aged host tissue to promote inflammaging.

### Exhausted GZMK+ T cells emerge in BALT with inflammaging

Aging in the lung is characterized by pro-inflammatory macrophage phenotypes concurrent with impaired lymphocyte function^12^, but it is not clear if this is associated with alterations of the resident immune architecture. Histologic analysis of aged (22-month) and young (2-month) wildtype (WT) mice (C57BL/6 strain) maintained in a specific-pathogen free (SPF) barrier facility demonstrated increased lymphoid aggregates adjacent to proximal airway and vessels previously described as inducible BALT^13^ in aged lungs (**Figures 1A and 1B**). BALT is a tertiary lymphoid structure that appears transiently in postnatal lungs^14^, but reemerges in adult disease states such as COPD^15^, autoimmunity^16^, and lung cancer^17^. To spatially resolve gene expression and cell types in BALT that develops in normally aged mouse lungs, we conducted spatial analysis of 429 unique mRNA probes using the Xenium platform that allowed clustering of major immune and structural cell types on Seurat with spatial coordinates (**Figure S1A**). Cellular composition analysis showed that BALTs in aged lungs are primarily composed of T and B cells alongside interstitial macrophages (IMs) and resident structural cells like fibroblasts (**Figures 1C and 1D**). Flow cytometry of resident immune cells (intravenous anti-CD45^−^) from aged lungs showed a relative increase in T (both CD4^+^ and CD8^+^) and B cells, along with increased interstitial macrophages (IMs) and a loss of alveolar macrophages (AMs) (**Figure S1B and S1C**). Flow cytometry of resident immune cells also demonstrated an increase in CD8^+^ T cells with exhaustion markers (PD-1^+^/TOX^+^) in aged lungs consistent with prior studies^9,18^ (**Figure S1B and S1C**), and spatial analysis demonstrated that markers of T cell exhaustion (including *Gzmk*) are primarily localized within BALTs in aged lungs (**Figure 1E**).

**Figure 1.**
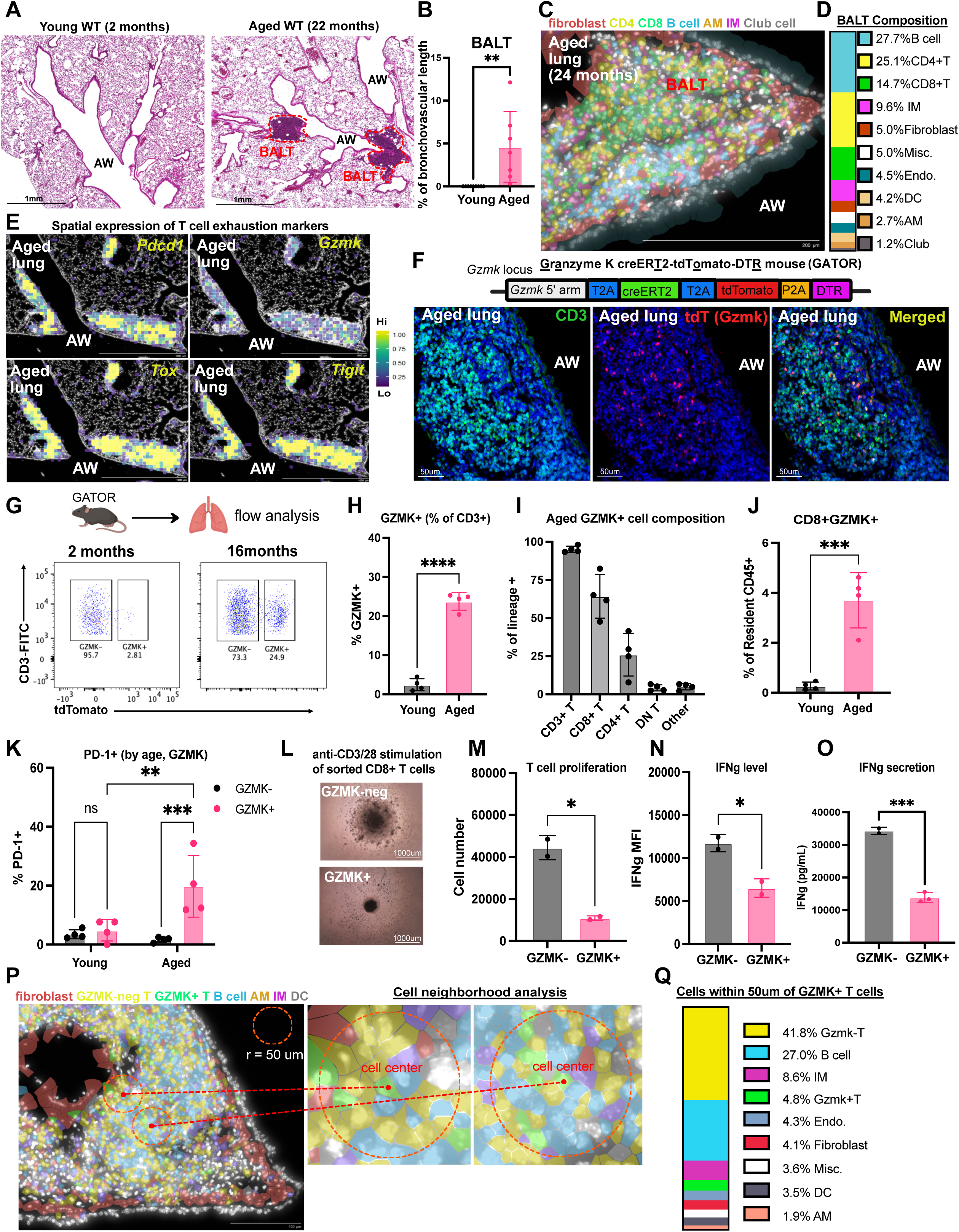
Exhausted GZMK+ T cells emerge in BALT with inflammaging. (A) H&E of lungs from young and aged wildtype B6 mice. (B) BALT quantification as percent length of airway and vessel with adjacent lymphoid tissue (n=9 young, n = 7 aged). (C-D) (C) Spatial analysis with Xenium platform on BALT from aged mouse along with (D) cellular composition (4 BALTs used for composition analysis). (E) Heatmaps of exhaustion genes in Xenium spatial analysis from aged lung. (F) Schematic of GATOR construct along with IHC of tdTomato+ (Gzmk+) CD3^+^ T cells within BALT of an aged GATOR mouse. (G-K) Flow cytometric analysis of aged vs young GATOR mice (n = 4 mice per group). (G-H) Gzmk+CD3+ lung resident T cells in GATOR mice by age. (I) Contribution of indicated cell types to total Gzmk^+^ lung resident cells from aged GATOR mice. (J) CD8^+^/Gzmk^+^ cells as a percentage of all resident CD45^+^ cells. (K) PD-1 expression among resident CD8^+^ T cells according to *Gzmk* expression and age. (L-O) Day 4 analysis of sorted resident GZMK^+^ and GZMK^−^/CD8^+^ T cells cultured with anti-CD3/CD28 beads (n = 2 wells per cell type). Cells incubated with brefeldin A for 4 hours prior to collection. (L) Brightfield imaging. (M) Total cell number and (N) mean fluorescence intensity (MFI) of IFNg among IFNg^+^ cells by flow cytometry. (O) IFNg ELISA of supernatant. (P-Q) (P) Method for spatial cell neighborhood analysis along with (Q) GZMK^+^ T cell neighborhood. All p-values determined by 2-tailed t-test except for panel k which was analyzed by 2-way ANOVA. * p<0.05, ** p<0.01, *** p<0.001, **** p<0.0001.

Recent studies have identified the emergence of GZMK^+^ T cells as a hallmark of inflammaging in aged tissues^9^. To characterize these T cells, we knocked in a polycistronic cassette that replaced the termination codon in the 3’ region of the *Gzmk* locus to generate the multifunctional (fluorescent reporter, lineage trace, cell deletion) GATOR mouse. Expression of *Gzmk* was minimal in young (2-month) lungs, but we could detect the emergence of *Gzmk*^+^ T cells in 16-month old GATOR mice by histology within BALTs (**Figures 1F and S1D**) and by flow cytometry (**Figures 1G, 1H, and S1E**). Lineage analysis of *Gzmk*^+^ cells demonstrated that over 90% are T cells (CD3^+^) composed primarily of CD8^+^ T cells (**Figures 1I and 1J**), and they co-express the exhaustion marker PD-1 with age (**Figure 1K**). Functional analyses demonstrated that *Gzmk*^+^/CD8^+^ T cells sorted from aged GATOR lungs failed to expand in response to T cell receptor (TCR) stimulation (anti-CD3/CD28 beads) (**Figures 1L and 1M**) and secreted lower levels of IFNγ relative to *Gzmk*^−^/CD8^+^ T (**Figures 1N, 1O and S1F**). To analyze the *Gzmk*^+^ T cell niche within tissues, we utilized our spatial data to import coordinates of cellular center points and identify nearest neighboring cells within a 50um radius of *Gzmk*^+^ T cells. Our “cell neighborhood” analysis demonstrated that *Gzmk*^+^ T are localized predominantly near T and B cells, but also IMs and structural cells such as endothelial cells and fibroblasts (**Figures 1P and 1Q)**. These data showed that exhausted *Gzmk*^+^ T cells emerge within organized tertiary lymphoid structures as part of the inflammaging phenotype.

### Age-related increase in NF-kB-activation in tissue fibroblasts

Emerging evidence supports an important role for the aging tissue microenvironment as a driver of age-related immune cell dysfunction^9,19^. We and others have shown that adventitial fibroblasts within the bronchovascular bundle serve as a niche for resident immune cells in the lung^20–22^, and spatial analysis demonstrated the presence of adventitial fibroblast markers expressed in and around BALT in aged lungs (**Figure 2A**). To evaluate aging hallmarks of adventitial fibroblasts, we examined the expression of *p16^Ink^*^4a^ in the lungs of young and aged INKBRITE (ultrasensitive *p16^Ink^*^4a–^GFP) reporter mice^23^. Flow cytometry demonstrated an increase in both the numbers of *p16^Ink^*^4a+^ adventitial fibroblasts and the absolute *p16^Ink^*^4a^ expression within adventitial fibroblasts in aged lungs (**Figures 2B and S2A**). *p16^Ink^*^4a+^ fibroblasts are primed for NF-kB activation in tissues^23,24^, and transcript analysis of sorted *p16^Ink^*^4a+^ adventitial fibroblasts from aged lungs demonstrated a significant enrichment of NF-kB target genes (relative to *p16^Ink^*^4a–^neg adventitial fibroblasts) (**Figure 2C**). We also performed single cell RNA-seq (scRNAseq) of lung fibroblasts isolated from young and aged lungs with clustering into previously annotated lung fibroblast subsets (adventitial, alveolar, and peribronchial)^25,26^ (**Figures 2D, S2B, and S2C**). Ingenuity pathway analysis (IPA) of aged vs. young fibroblast subsets showed that differentially expressed genes (DEG) enriched in adventitial fibroblasts were highly enriched for pathways activating NF-kB (**Figures 2E, S2D and Supplementary Table 1**), and expression analysis showed upregulation of NF-kB targets in fibroblasts (**Figures 2F and S2E**) similar to those found upregulated specifically in aged *p16^Ink^*^4a+^ adventitial fibroblasts (**Figure 2C**). Immunocytochemistry analysis demonstrated increased p65/RelA nuclear localization in cultured aged adventitial fibroblasts (**Figure 2G)**, confirming elevated NF-kB activation.

**Figure 2.**
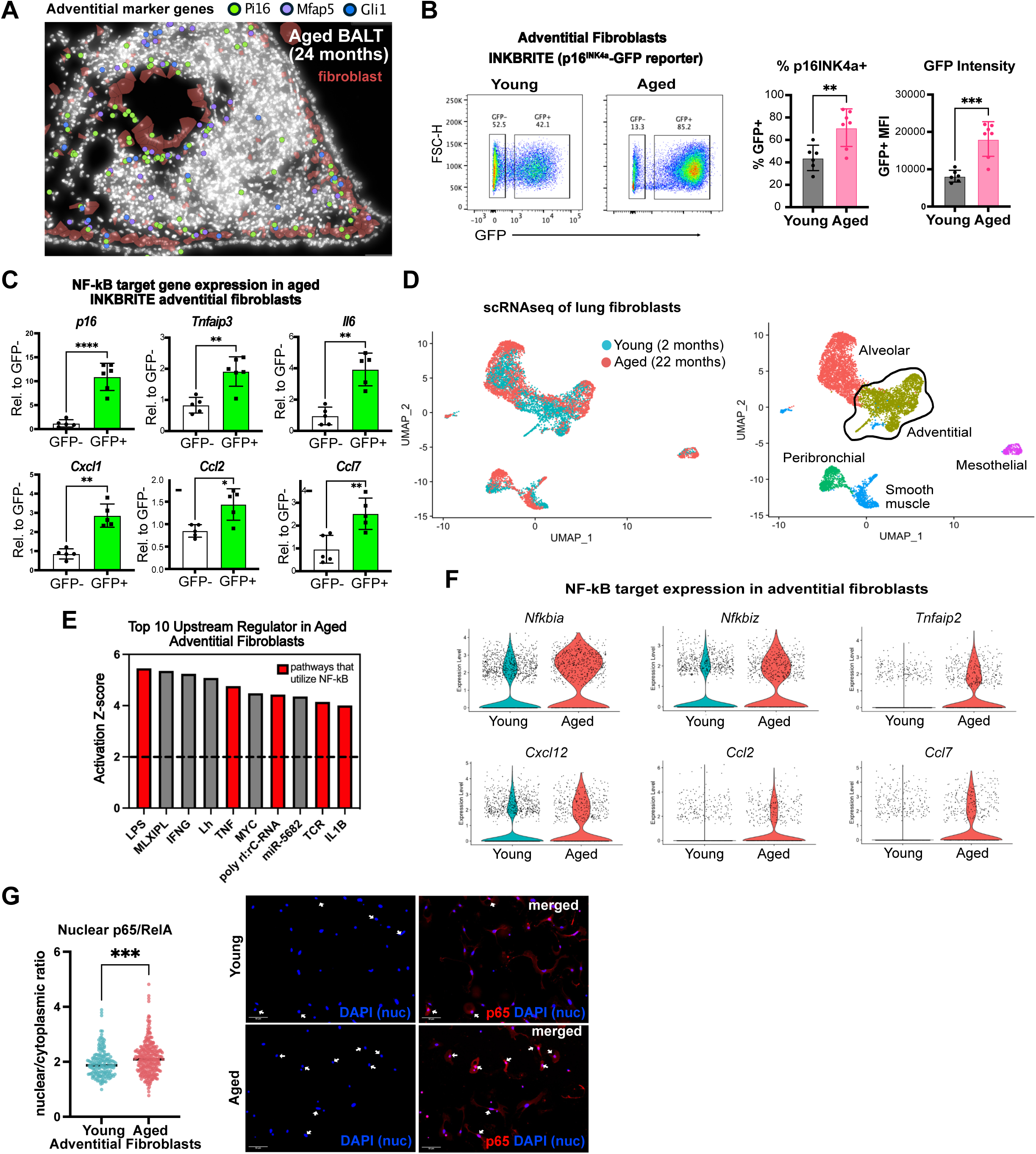
Age-related increase in NF-kB-activation in tissue fibroblasts. (A) Xenium spatial analysis of BALT from aged mouse identifying adventitial fibroblast markers. (B) Flow cytometry plots of GFP expression in adventitial fibroblasts from young (4-month) and aged (24-month) INKBRITE mice, as well as percentage of fibroblasts that are GFP+, and GFP intensity within GFP+ population (young n = 6 mice, aged n = 7 mice). (C) qPCR analysis of indicated NF-kB target genes in GFP+ vs GFP-adventitial fibroblasts from aged (24-month) INKBRITE mice. (D) UMAPs of scRNA-sequencing data of fibroblasts from young and aged WT mice. (E) IPA of scRNA-sequencing of aged vs. young adventitial fibroblasts. Activation score >2 predicts pathway activation. (F) Violin plots of NF-kB target genes in adventitial fibroblasts. (G) Nuclear/cytoplasmic ratio of p65 in cultured lung adventitial fibroblasts from young (3 mo.) versus aged (>24 mo.) WT mice, as well as sample anti-p65 immunocytochemistry. Ratio determined by intensity of p65 staining overlying the nucleus compared with that of the cytoplasm (young n = 171, aged n = 279 cells). All p-values determined by 2-tailed t-test, * p<0.05, **p<0.01, *** p<0.001, ****p<0.0001.

### NF-kB-activated fibroblasts promote the inflammaging phenotype

NF-kB initiates negative feedback loops by activating transcription of genes encoding negative regulators^27^ such as *Tnfaip3* and *Nfkbia* in aged adventitial fibroblasts (**Figure 2F**). TNFAIP3 is a ubiquitin-modifying enzyme that prevents nuclear localization of the NF-kB complex, the loss of which de-represses NF-kB activation^28^ (**Figure 3A**). To assess whether NF-kB activation in fibroblasts contribute to the aging phenotype, we deleted *Tnfaip3* utilizing a pan-fibroblast Cre mouse^29^ (*Dermo1^Cre/+^:Tnfaip3^fl/fl^*, or *Dermo1^Tnfaip^*^3^*^−CKO^*) (**Figure S3A**). By 3 months of age, *Dermo1^Tnfaip^*^3^*^−CKO^* mice demonstrated spontaneous formation of BALTs (**Figures 3B and 3C**). To evaluate changes in the lung resident immune compartment we performed scRNAseq of tissue resident immune cells from *Dermo1^Tnfaip^*^3^*^−CKO^* and controls, which demonstrated high enrichment of CD8^+^ T cells in the *Dermo1^Tnfaip^*^3^*^−CKO^* lungs (**Figures 3D, 3E and S3B**).

**Figure 3.**
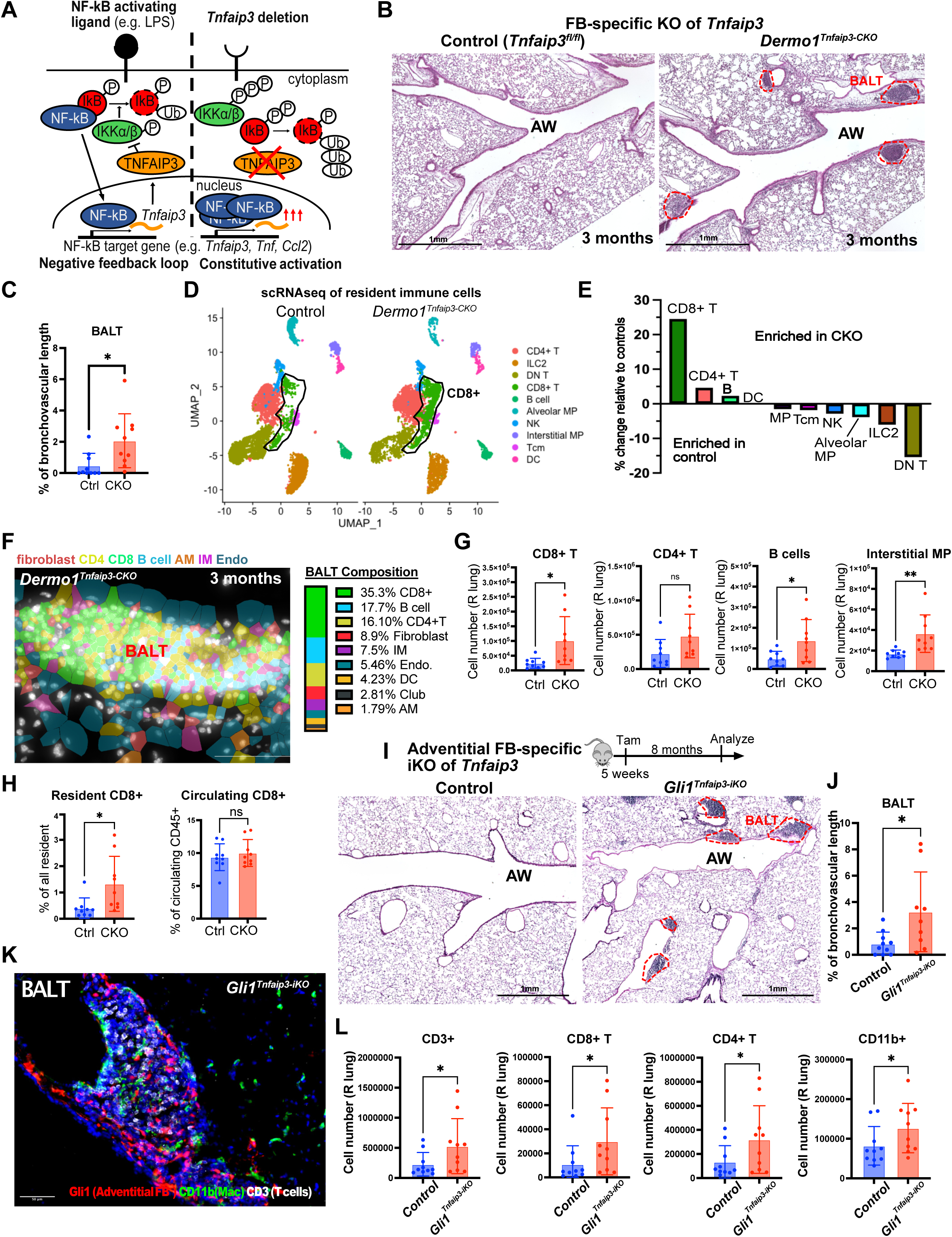
NF-kB-activated fibroblasts promote the inflammaging phenotype. (A) Schematic of how deletion of *Tnfaip3* leads to increased NF-kB activation. (B) H&E of lung sections from young control and fibroblast conditional knock-out (CKO) of *Tnfaip3* (*Dermo1^Tnfaip^*^3^*^−CKO^*) mice. (C) BALT as percent of total length of airways and vessels with adjacent lymphoid tissue (n=10 mice per genotype). (D) UMAP of scRNA-sequencing of lung resident CD45+ cells from 3-month-old *Tnfaip3f/f* (Control) and *Tnfaip3* CKO mice according to genotype. (E) Percent change in composition of immune cell subsets among total resident CD45+ cells between Control and *Tnfaip3* CKO mice according to scRNA-seq. (F) Cellular composition of BALT from *Dermo1^Tnfaip^*^3^*^−CKO^* mouse according to Xenium spatial transcriptome analysis (4 BALTs used for composition analysis). (G-H) Flow cytometric analysis of immune cells from 3-month-old *Tnfaip3f/f* (Ctrl) or *Dermo1^Tnfaip^*^3^*^−CKO^* (CKO) mice (n = 9 mice per group). (G) Lung resident immune cell numbers. (H) CD8+ T cells as a fraction of total lung resident cells (ivCD45-) or circulating (ivCD45+) cells. (I – L) Analysis of control (*Tnfaip3f/f*) and inducible adventitial fibroblast knock-out (iKO) of *Tnfaip3* (*Gli1^Tnfaip^*^3^*^−iKO^*) mice 8 months after *Tnfaip3* deletion (n=9-10 mice per genotype). (I) H&E staining and (J) BALT quantification by genotype. (K) IHC of BALT from *Gli1^Tnfaip^*^3^*^−iKO^* mouse. (L) Flow cytometry of lung resident immune cells from control and *Gli1^Tnfaip^*^3^*^−iKO^* mice. All p-values determined by 2-tailed t-test, except for panel (L), which was analyzed by 1-tailed t-test. * p<0.05, *** p<0.001.

Spatial analysis of the BALTs in 3-month-old *Dermo1^Tnfaip^*^3^*^−CKO^* lungs was consistent with scRNAseq findings and identified T and B cell predominant lymphoid clusters that also had a sizeable fraction of fibroblasts and IMs (**Figures 3F and S3C**). Findings were confirmed with flow cytometry, which demonstrated increased tissue resident (intravenous anti-CD45^−^) T (particularly CD8^+^ T cells) and B cells along with IMs compared to controls (**Figures 3G and S3D**) that mimicked 22-month-old WTs.

Importantly, we did not observe a similar increase in the fraction of circulating CD8+ T cells, demonstrating the importance of the lung microenvironment in driving CD8+ T cell accumulation (**Figure 3H**). We also activated NF-kB in the lung epithelium by deleting *Tnfaip3* in a lung epithelium-specific Cre mouse^30^ (*Nkx2.1^Cre/+^:Tnfaip3^fl/fl^*), which failed to generate BALT or increase resident immune cells in 3-month-old lungs (**Figures S4A-C**). To determine whether inducible activation of NF-kB specifically in adventitial fibroblasts was sufficient to promote the inflammaging phenotype, we utilized an adventitial fibroblast creERT2 mouse^31^ (**Figure S4D**) to delete *Tnfaip3* (*Gli1^CreERT^*^2^*^/+^:Tnfaip3^fl/fl^*or *Gli1^Tnfaip^*^3^*^−iKO^*). Tamoxifen was administered to *Gli1^Tnfaip^*^3^*^−iKO^* and controls at 5 weeks of age followed by an 8-month chase period prior to analysis.

Histologic analysis demonstrated an increase in BALT formation in *Gli1^Tnfaip^*^3^*^−iKO^* animals (**Figures 3I and 3J)**, and lineage trace of the *Gli1*^+^ fibroblasts (with R26^tdTomato^) in the *Gli1^Tnfaip^*^3^*^−iKO^* animals demonstrated infiltration of adventitial fibroblasts within the BALTs formed after deletion of *Tnfaip3* (**Figure 3K**). Flow cytometry showed an increase in resident T (CD4^+^ and CD8^+^) cells along with IMs (**Figures 3L and S4E**). These results demonstrated that NF-kB activation within a resident fibroblast subset is sufficient to accelerate the formation of tertiary lymphoid structures seen in normal aging.

### Fibroblastic NF-kB activation induces CD8^+^ T cell clonal expansion and exhaustion

To define the topography of *Gzmk*^+^ T cells in our accelerated model of inflammaging, we generated *Dermo1^Tnfaip^*^3^*^−CKO^*:GATOR mice and performed thick section imaging of 3-month old animals. Imaging demonstrated resident (IV CD45^−^) *Gzmk*^+^ T cells (tdTomato^+^/CD3^+^) accumulating predominantly within lymphoid aggregates along the airway (marked by aSMA) (**Figure 4A**). Spatial gene expression analysis of *Dermo1^Tnfaip^*^3^*^−CKO^*lungs confirmed the localization of *Gzmk* transcripts along with other markers of T cell exhaustion (*Pdcd1, Tox, Tigit*) within BALTs (**Figure 4B**), resembling WT aged lungs. Segregation of CD8^+^ T cells by *Gzmk* expression on thick section imaging showed that *Gzmk*^+^/CD8^+^ T cells appear spatially clustered together (**Figure 4C**).

**Figure 4.**
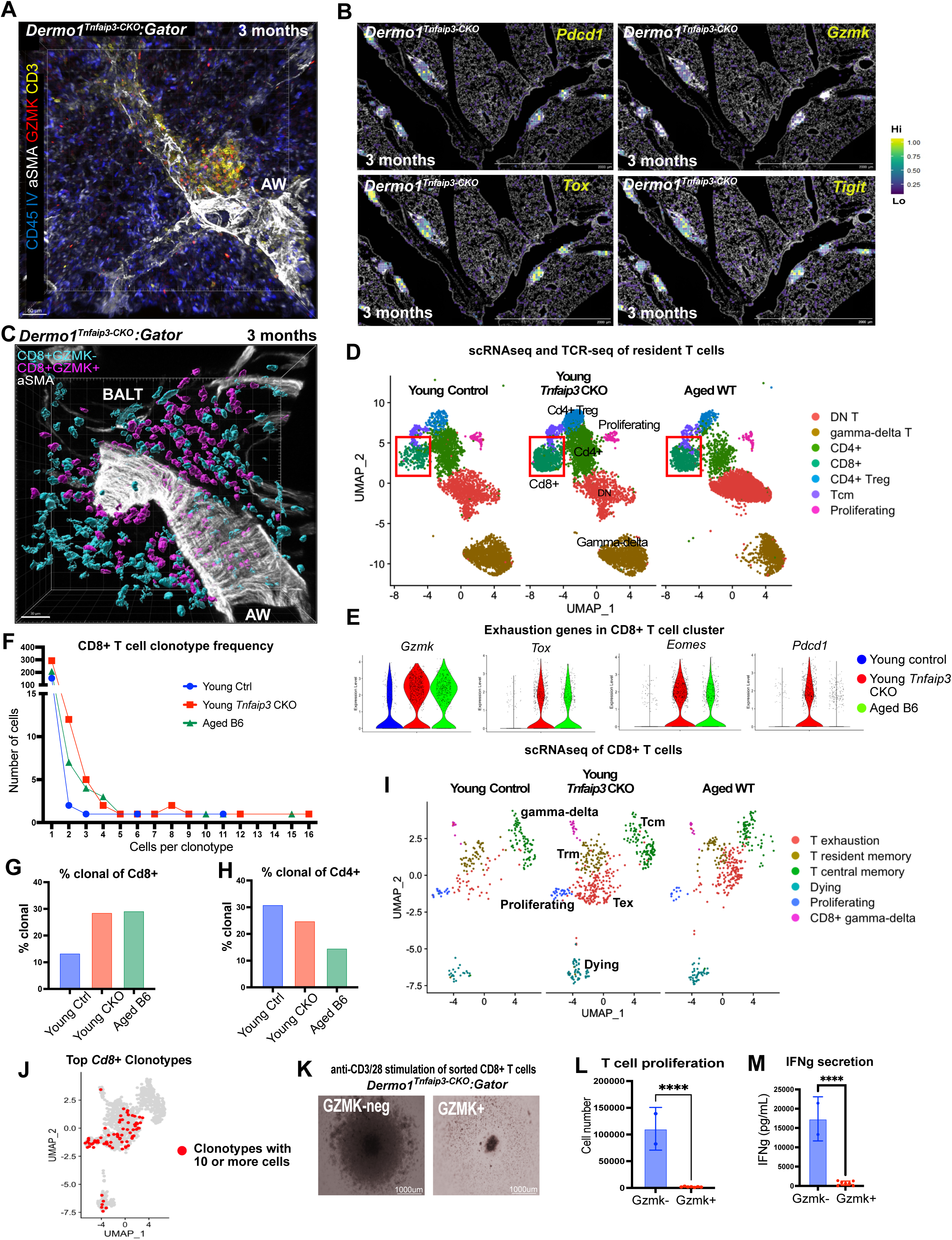
Fibroblast NF-kB activation induces CD8+ T cell clonal expansion and exhaustion. (A) Thick section imaging of lung with BALT from a *Dermo1^Tnfaip^*^3^*^−CKO^:GATOR* mouse. (B) Heatmaps of exhaustion genes in Xenium spatial analysis from *Dermo1^Tnfaip^*^3^*^−CKO^* lung. (C) Thick section imaging of *Dermo1^Tnfaip^*^3^*^−CKO^* mouse demonstrating localization of GZMK^+^ and GZMK^−^/CD8^+^ T cells within a BALT. (D) UMAPs of scRNA-sequencing of lung resident CD3^+^ T cells from 3-month-old *Tnfaip3f/f* (Young control), 3-month-old *Dermo1^Tnfaip^*^3^*^−CKO^* (Young *Tnfaip3* CKO) and 23 month (aged WT) mice. (E) Violin plots of T cell exhaustion genes within CD8^+^ T cell cluster. (F) Number of sequenced *Cd8^+^* T cells with shared TCR (clonotype) by sample. (G-H) Percent of total lung resident (G) *Cd8*^+^ or (H) *Cd4*^+^ T cells that are clonal (more than one cell with shared TCR). (I) UMAP of sub-clustered *Cd8*^+^ T cells by sample and cell type. (J) UMAP identifying all Cd8^+^ T cells from which the same TCR was identified in 10 or more cells. (K-M) Day 4 analysis of lung resident GZMK^+^ and GZMK^−^/CD8^+^ T cells sorted from *Dermo1^Tnfaip^*^3^*^−CKO^;GATOR* mice and cultured for with anti-CD3/CD28 beads (Gzmk-neg n= 2 wells, Gzmk-pos = 7 wells). (K) Brightfield imaging. (L) Total cell number as determined by flow cytometry. (M) IFNg ELISA of supernatant. p-values determined by 2-tailed t-test. **** p<0.0001.

To compare the transcriptome of T cells arising in normal aging and our accelerated inflammaging model, we performed single cell TCR sequencing coupled with scRNAseq on tissue-resident T cells sorted from young (3 months) control and *Dermo1^Tnfaip^*^3^*^−CKO^*lungs along with WT aged (23 months) lungs. Cluster analysis with uniform manifold approximation and projection (UMAP) showed that *Dermo1^Tnfaip^*^3^*^−CKO^* and aged lungs are enriched for *Gzmk*^+^/CD8^+^ T cells (**Figures 4D, 4E and S5A-C**). TCR analysis of *Cd8*+ T cells revealed a similar increase in clonality among *Dermo1^Tnfaip3CKO^* and aged WT mice compared with young controls, but not within the *Cd4+* T cell population (**Figures 4F-H**). Re-clustering of the *Cd8*+ cells revealed a distinct population of cells with features of exhaustion in the *Dermo1^Tnfaip3CKO^* and aged WT mice that were minimally represented in young controls (**Figures 4I, S5D-F, and Supplementary Table 2**). Integration of TCR clonotypes with single cell transcriptome showed that the expanded CD8^+^ T cell clones include proliferating, exhausted, and dying T cells (**Figures 4J and S5G)** suggesting that exhausted state is accompanied by initial proliferative activation. Flow cytometry of *Dermo1^Tnfaip^*^3^*^−CKO^*and *Gli1^Tnfaip^*^3^*^−iKO^*lungs demonstrated an increase in PD-1^+^/TOX1^+^/CD8^+^ T cells relative to controls (**Figures S5H and S5I**). PD-1^+^/TOX1^+^/CD8^+^ T cells in *Dermo1^Tnfaip^*^3^*^−CKO^*lungs exhibited lower levels of IFNγ relative to PD-1^−^/TOX1^−^/CD8^+^ T cells (**Figure S5J**). Additionally, *Gzmk*^+^/CD8^+^ T cells sorted from young *Dermo1^Tnfaip^*^3^*^−CKO^*:GATOR mice lungs failed to expand in response to TCR stimulation and secreted lower levels of IFNγ relative to *Gzmk*^−^/CD8^+^ T cells (**Figures 4K-M**). These results demonstrate that fibroblastic NF-kB activation induces polyclonal expansion and functional exhaustion of tissue resident CD8^+^ T cells comparable to what had been reported in aged organs^9,32^.

### A fibroblast-macrophage-T cell circuit promotes the emergence of GZMK^+^ T cells

We profiled the bulk transcriptome of fibroblasts sorted from the lungs of young *Dermo1^Tnfaip^*^3^*^−CKO^*and controls. Differentially expressed gene (DEG) analysis showed significant (p < 0.05) upregulation of NF-kB targets in the *Dermo1^Tnfaip^*^3^*^−CKO^* samples (**Figure 5A and Supplementary Table 3**). Hypergeometric probability testing of upregulated DEGs from fibroblasts of *Dermo1^Tnfaip^*^3^*^−CKO^* demonstrated significant overlap with genes enriched in aged adventitial fibroblasts (but not aged alveolar fibroblasts) (**Figures 5B and S6A**). Upregulated genes shared between *Dermo1^Tnfaip^*^3^*^−CKO^*and aged adventitial fibroblasts include genes involved in antigen presentation (*B2m, H2-D1, H2-K1*, etc.), immune cell recruitment (*Ccl2, Ccl7, Ccl19*) and complements (*C3, C4b*). To determine whether NF-kB activated fibroblasts directly promote T cell expansion and exhaustion *in vitro*, we performed fibroblast-T cell coculture with lung fibroblasts isolated from *Tnfaip3^fl/fl^* animals treated with adeno-Cre or adeno-empty vectors. Downregulation of *Tnfaip3* in fibroblasts (adeno-Cre deletion) induced NF-kB target genes (**Figure S6B**) but failed to induce expansion or exhaustion markers in splenic T cells when cocultured together (**Figure 5C**). *Tnfaip3*-deleted fibroblasts also failed to increase T cell recruitment in a transwell migration assay (**Figure 5D**), which suggested that NF-kB activated fibroblasts do not act directly on T cells to induce expansion, exhaustion, or recruitment.

**Figure 5.**
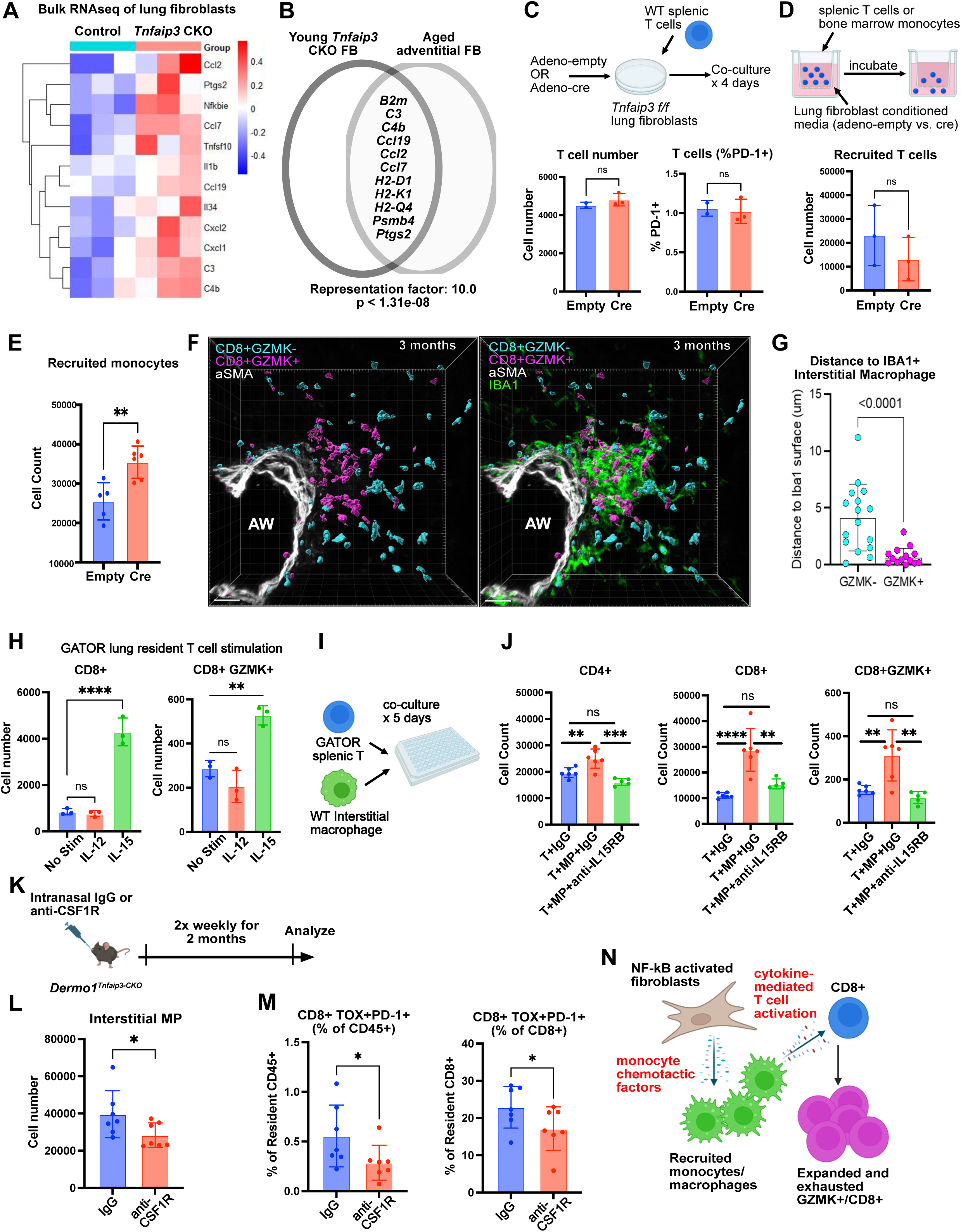
A fibroblast-macrophage-T cell circuit promotes the emergence of GZMK+ T cells. (A) Heatmap of NF-kB target genes in lung fibroblasts from young (2 mo.) control (*Tnfaip3f/f*) and *Dermo1^Tnfaip^*^3^*^−CKO^* (*Tnfaip3* CKO) mice. Each column = biological replicate. (B) Genes upregulated in both fibroblasts from young *Tnfaip3* CKOs vs. controls, and aged (22 mo.) vs. young (2 mo.) adventitial fibroblasts. Representation factor > 1 indicates more overlap than expected between two independent groups. (C) Flow cytometric analysis of T cells after co-culture with fibroblasts with or without deletion of *Tnfaip3* (2-3 wells per condition). (D-E) Transwell migration assay with (D) T cells (3 wells per condition) or (E) monocytes (5-6 wells per condition) incubated with conditioned media from fibroblasts with or without deletion of *Tnfaip3*. (F-G) (F) Thick section imaging of BALT from *Dermo1^Tnfaip^*^3^*^−CKO^:*GATOR mice and (G) distance between GZMK^+^/CD8^+^ vs GZMK^−^/CD8^+^ T cells and IBA1^+^ interstitial macrophages (IM) (n = 5 mice, 16 clusters). (H) Day 5 analysis of lung resident T cells incubated with indicated cytokines. (I-J) (I) Experimental model and (J) day 5 analysis of T cells cultured alone or with IM and indicated antibodies. (K) Experimental model for depletion of IMs. (L) IM number in IgG or anti-CSF1R treated mice. (M) PD-1^+^/TOX1^+^/CD8^+^ cells after 2 months of antibody treatment (n=7 mice per group). (N) Proposed mechanism by which lung fibroblast NF-kB activation induces the expansion and exhaustion of CD8+ T cells. Panels C-E and G analyzed by 2-tailed t-test, H and J by 1-way ANOVA, and (L-M) by 1-tailed t-test. * p<0.05, ** p<0.01, *** p<0.001, **** p<0.0001.

Fibroblasts have been shown to recruit IMs derived from circulating monocytes^33,34^. IMs constitute a sizeable fraction of the immune cells in BALTs and are increased in both WT aged and *Dermo1^Tnfaip^*^3^*^−CKO^*lungs (**Figures 1, 3**). Conditioned media from *Tnfaip3*-deleted fibroblasts significantly enhanced monocyte recruitment across transwells compared to *Tnfaip3*-intact fibroblasts (**Figure 5E**). Topographic analysis of IBA1+ IMs within BALTs from *Dermo1^Tnfaip^*^3^*^−CKO^*:GATOR mice demonstrated that *Gzmk*^+^/CD8^+^ T cells are situated closer to IBA1+ IMs than *Gzmk*^−^/CD8^+^ T cells (**Figures 5F and 5G**), which would enable intercellular interactions. Myeloid cell-derived cytokines such as IL-15 and IL-12 had been reported to induce T cell exhaustion markers independent of TCR stimulation^35^, and they are also upregulated in the IMs of *Dermo1^Tnfaip^*^3^*^−CKO^* lungs (**Figure S6C**). Stimulation of lung-resident (IV CD45^−^) or circulating T cells isolated from GATOR demonstrated that IL-15 but not IL-12b significantly increased CD8^+^ T cell expansion and induction of *Gzmk* expression in the absence of TCR stimulation (**Figures 5H and S6D**). Coculture of splenic T cells from GATOR with lung IMs also induced CD8^+^ T cell expansion and *Gzmk* induction, which was blocked in the presence of neutralizing antibody to IL-15Rb (**Figures 5I and 5J**). Finally, to test the effect of macrophage depletion on exhausted T cells *in vivo*, we administered anti-CSF1R or isotype control to *Dermo1^Tnfaip^*^3^*^−CKO^*animals (**Figure 5K**). Anti-CSF1R reduced IMs in the lung concurrent with a decrease in tissue resident PD-1^+^/TOX1^+^/CD8^+^ T cells (**Figures 5L, 5M and S6E**). These data illustrate a fibroblast-macrophage-T cell circuit triggered by fibroblastic NF-kB activation that promotes IM recruitment, which in turn drives CD8+ T cell expansion and exhaustion (**Figure 5N**) within tertiary lymphoid structures in inflammaging.

### GZMK+ T cells promote inflammatory response in ARDS

Neutrophil influx into the lung is a defining hallmark of ARDS^36^, and neutrophil count is directly correlated with ARDS mortality, whereas resident AMs (but not IMs) were found to have a protective role^37^. Even prior to COVID-19, age was identified as one of the strongest risk factors for the development of ARDS that follows common infections or trauma^38^. In a model of ARDS induced by intranasal administration of lipopolysaccharide (LPS), aged (18-month old) mice developed an exaggerated ARDS phenotype compared to young animals, as demonstrated by an increase in immune infiltration in the lung, along with increased neutrophils in the extravascular space (IV CD45^−^) and loss of AMs (**Figures 6A, 6B, S7A and S7B**). We induced ARDS in our accelerated inflammaging model (*Dermo1^Tnfaip^*^3^*^−CKO^*) at 2 months of age. Like aged WT mice, *Dermo1^Tnfaip^*^3^*^−CKO^*animals demonstrated exaggerated immune cell infiltration that is primarily driven by increased neutrophilic influx compared to controls, with a non-significant trend (p = 0.059) towards reduction in AMs (**Figures 6C, 6D and S7C**). The difference in inflammatory infiltrates was largely driven by females in the cohort (**Figure S7D**), consistent with prior report of increased inflammatory response to LPS in female mice^39^. Cytokine quantification of lung homogenates demonstrated elevation of proinflammatory mediators, including the neutrophil chemoattractants CXCL1 and CXCL2 in the *Dermo1^Tnfaip^*^3^*^−CKO^* samples (**Figures 6E and S7E**). Histologic evaluation of *Dermo1^Tnfaip^*^3^*^−CKO^* lungs demonstrated significantly higher grades of alveolar damage as measured by the standardized Lung Injury Scoring System^40^ (**Figures 6F and 6G**). These results demonstrated that our accelerated inflammaging model recapitulated the increased susceptibility to ARDS seen in aged WT.

**Figure 6.**
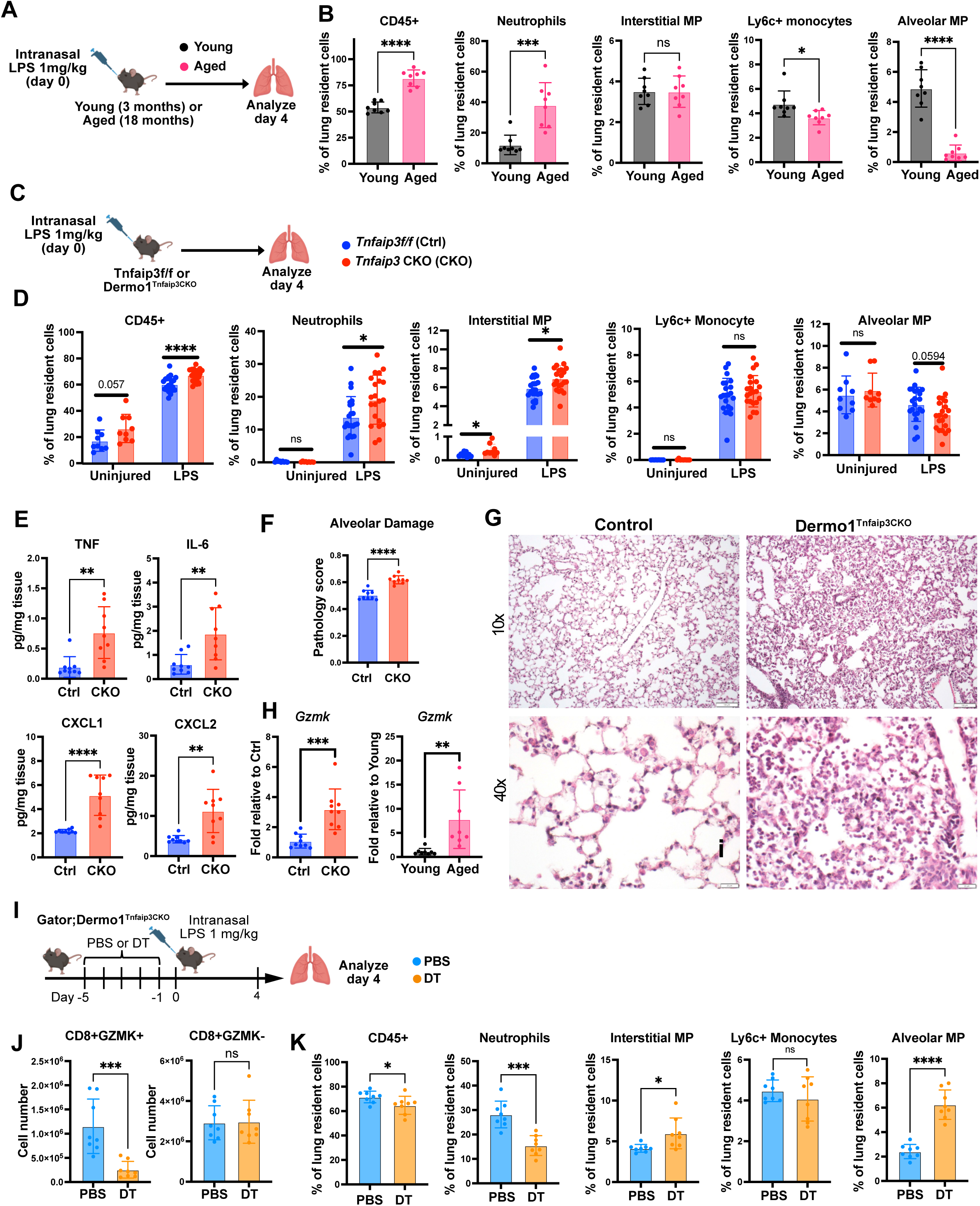
GZMK+ T cells promote inflammatory response in ARDS. (A-B) Young or aged WT (C57BL/6) mice were administered intranasal lipopolysaccharide (LPS) and analyzed on day 4 (n = 8 mice per group). (A) Experimental model and (B) Immune populations as a percentage of total lung resident cells. (C-G) 3-month control or *Tnfaip3* CKO mice were administered intranasal LPS and analyzed on day 4. (C) Experimental model. (D) Immune populations as a percentage of total lung resident cells (n = 21-22 per group). Immune populations in homeostasis from a similarly age and sex-matched cohort included for comparison (n = 9 mice per group). (E) Protein concentrations from lung homogenates (n = 9 mice per group). (F) Quantification of alveolar damage of control and *Tnfaip3* CKO mice (n = 9 mice per group) and (G) example H&E staining. (H) qPCR of *Gzmk* expression in whole lung homogenates from indicated cohorts on day 4 post-LPS (n = 8-9 mice per group). (I-K) Analysis of *Dermo1^Tnfaip^*^3^*^−CKO^;*GATOR mice day 4 post LPS with and without prior GZMK cell depletion (n = 8 mice per group). (I) Experimental schematic for *Gzmk*^+^ T cell depletion prior to intranasal LPS. (J) Number of lung resident GZMK^+^ and GZMK^−^/CD8^+^ T cells. (K) Immune populations as a percentage of total lung resident cells. Statistical analyses done by 2-tailed t-test. * p<0.05, ** p<0.01, *** p<0.001, **** p<0.0001.

While exhausted T cells emerge during inflammatory states, they are often thought to protect tissue against excessive inflammation by restraining T cell activation. GZMK, unlike GZMA, exhibits limited cytolytic activity^10,41^, but has been found to augment inflammatory output in fibroblasts^9,10,42^. We stimulated lung fibroblasts with recombinant GZMK and found mild increases in inflammatory mediators (**Figure S8A**). However, GZMK had a more potent effect on macrophages *in vitro*, with larger fold inductions of chemotactic factors that mediate neutrophil recruitment (e.g. *Cxcl1* and *Cxcl2*) (**Figure S8B**). Lungs from both aged WT and *Dermo1^Tnfaip^*^3^*^−CKO^* animals exhibited elevated *Gzmk* expression day 4 post-LPS (**Figure 6H**) concordant with the finding of elevated *Gzmk*^+^ T cells in these lungs. To characterize the effect of *Gzmk*^+^ T cells on ARDS *in vivo*, we administered diptheria toxin (DT) or PBS to *Dermo1^Tnfaip^*^3^*^−CKO^*:GATOR mice prior to induction of ARDS (**Figure 6I**). DT administration efficiently removed *Gzmk*^+^/CD8^+^ T cells in the lung without altering *Gzmk*^−^/CD8^+^ T cells (**Figure 6J**). Deletion of *Gzmk*^+^ cells resulted in a dramatic reduction of neutrophilic influx into the lung, while preserving the number of AMs (**Figures 6K, S8C and S8D**). These results demonstrated that exhausted *Gzmk*+ T cells triggered by fibroblastic NF-kB activation can drive underlying susceptibility to inflammation in tissues with the inflammaging phenotype.

## Discussion

Aging increases susceptibility to adult-onset lung diseases, including ARDS, emphysema and pulmonary fibrosis^43^. These diseases have all been linked with inflammaging, yet specifically how age-associated immune dysregulation predisposes individuals to disease remains largely unknown. At the same time, it is becoming clear that our immune system is not aging in isolation, and the aging tissue microenvironment can impact immune cell function^19,44^. In this study, we showed that fibroblasts orchestrate the structural remodeling of the pulmonary immune system associated with inflammaging through NF-kB activation (**Figure S9**). A hallmark of fibroblast aging is the acquisition of senescence and inflammatory states^45^, both of which have been associated with NF-kB activation^24,46^. NF-kB has been shown to program fibroblast states in pathologic settings such as a cancer^47,48^, but here we show that it is also part of an aging program to remodel the immune architecture. We show that NF-kB triggers a fibroblast-macrophage-T cell circuit that leads to polyclonal expansion of an exhausted *Gzmk*^+^/CD8^+^ T cell population within BALT. This is analogous to what was described in lymph nodes with fibroblastic reticular cells coordinating the recruitment and maintenance of dendritic cells required for T cell priming^49,50^, suggesting a conserved fibroblast-myeloid cell-lymphocyte circuit that influences adaptive immunity in peripheral tissues. Our work provides a link between a hallmark of fibroblast aging with another hallmark of immune cell aging, underscoring how aging structural cells shape the local microenvironment to re-organize the resident immune architecture.

Another intriguing finding from this study was that activation of fibroblast NF-kB led to the clonal expansion and exhaustion of CD8^+^ T cells without the introduction of a chronic antigen. Our mice are maintained in SPF conditions but not germ-free, so we could not exclude the presence of endogenous microbiome that may drive the exhaustion phenotype through TCR stimulation. However, we do show that macrophages induced the expansion and induction of *Gzmk*+ T cells in the absence of TCR stimulation *in vitro*. This data is consistent with prior work showing that monocyte/macrophage-derived cytokines can activate memory T cells in an antigen-independent fashion^51,52^. Tumor-associated macrophages have been shown to prime T cells for exhaustion in the tumor microenvironment through antigen-presentation^53,54^, but this does not exclude their ability to promote exhaustion in bystander T cells. While T cell exhaustion has been broadly defined as a response to chronic antigen stimulation, we show that the *Gzmk^+^*/CD8^+^ T cells induced in response to fibroblastic NF-kB activation bear many of the hallmarks of exhausted T cells (expression of inhibitory receptors, decreased cytokine production, reduced proliferative capacity in response to TCR stimulation, association with clonal expansion, and etc.)^55^. It will be important to determine in future studies whether all or only particular subsets of T cells are susceptible to this cytokine driven exhaustion, and whether T cells that develop features of exhaustion in an antigen-independent manner are functionally distinct from those that develop through chronic antigen stimulation.

Despite growing literature that has identified the presence of *Gzmk^+^*/CD8^+^ T cells in multiple disease contexts^10,11,42^, defining the function of these cells in pathogenesis has been challenged by the lack of adequate tools. Utilizing our novel GATOR mouse, we demonstrated that *Gzmk*^+^ T cells are not merely byproducts of inflammaging, but rather a potent driver of tissue inflammation and modifier of age-related inflammatory response. GZMK appears to have a role in stimulating myeloid cell recruitment to tissues, which could lead to further NF-kB activation in the stroma and induction of T cell exhaustion in a positive feedback loop to maintain the inflammaging phenotype. Additionally, our finding that both aged adventitial fibroblasts and NF-kB activated fibroblasts upregulate complement proteins is also of significance given multiple recent studies that identified complement activation as a critical function of GZMK^56,57^, providing a mechanism by which aged tissues may be more susceptible to GZMK-induced inflammation. Together, these findings demonstrate that at least in certain contexts, T cell exhaustion can simultaneously limit adaptive immunity while augmenting the innate immune response, which is one of the hallmarks of inflammaging. These results suggest that *Gzmk*^+^ T cells are potential targets for rejuvenation in aged tissues to modify their susceptibility to age-related diseases.

## Author Contributions

N.C.A. and T.P. conceived the experiments and wrote the manuscript. N.C.A., C.R., J.Y.L., N.R., R.B., M.M., P.R., V.A. performed the experiments, collected samples, and analyzed data. M.L., A.M., A.B.M provided input on the experiments and manuscript.

## Acknowledgements

We thank Parnassus Flow Cytometry Core, and in particular Vinh Nguyen, for assistance with cell sorting (P30DK063720) and Walter Eckalbar at the Genomics CoLab at UCSF for assistance in processing single cell and TCR sequencing data. This work is supported by NIH grants R01HL160895 and Bakar Aging Research Institute (BARI) research award to T.P., BARI postdoctoral award, F32HL156452, and K08HL169723 to N.C.A.

---

**Figure S1.**
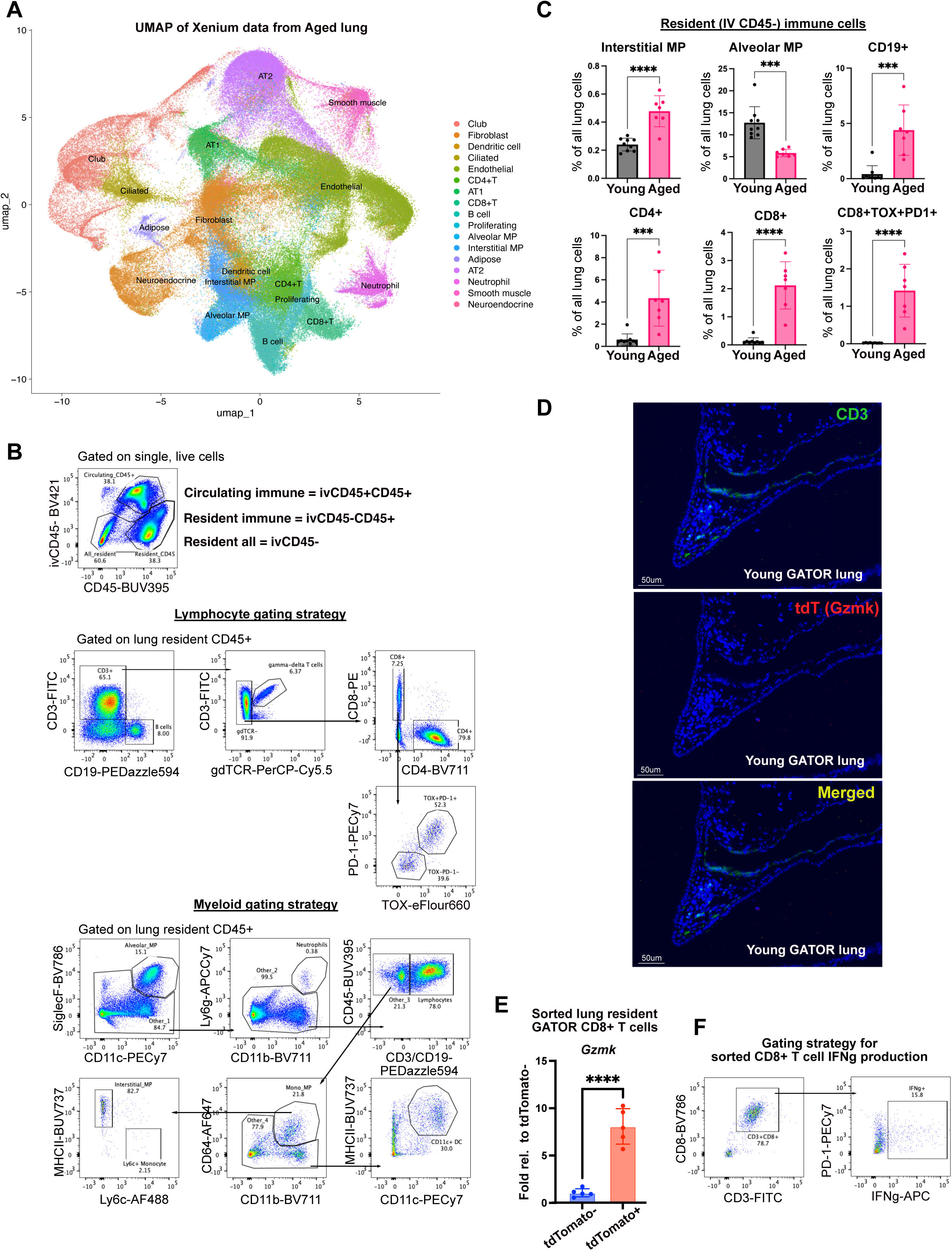
Characterization of the changing lung immune landscape with age. (A) Seurat clustering and UMAP from Xenium spatial transcriptome analysis of aged (24 mo.) lung. (B) Flow cytometry gating strategy to identify resident versus circulating immune cells as well as lymphocyte and myeloid subpopulations. (C) Flow cytometric analysis of resident immune cells from young (2 month, n = 9 mice) and aged (22 month, n = 7 mice) WT C57BL/6 mice as a percentage of all lung cells (ivCD45+ excluded). (D) IHC of CD3+ T cells in young (2-month) GATOR mice. (E) qPCR of *Gzmk* expression in sorted lung resident tdTomato-vs + CD8+ T cells from GATOR mice (n = 5 mice). (F) Flow cytometry gating strategy for quantifying IFNg level. Statistical analyses done by 2-tailed t-test. * p<0.05, ** p<0.01, *** p<0.001, **** p<0.0001.

**Figure S2.**
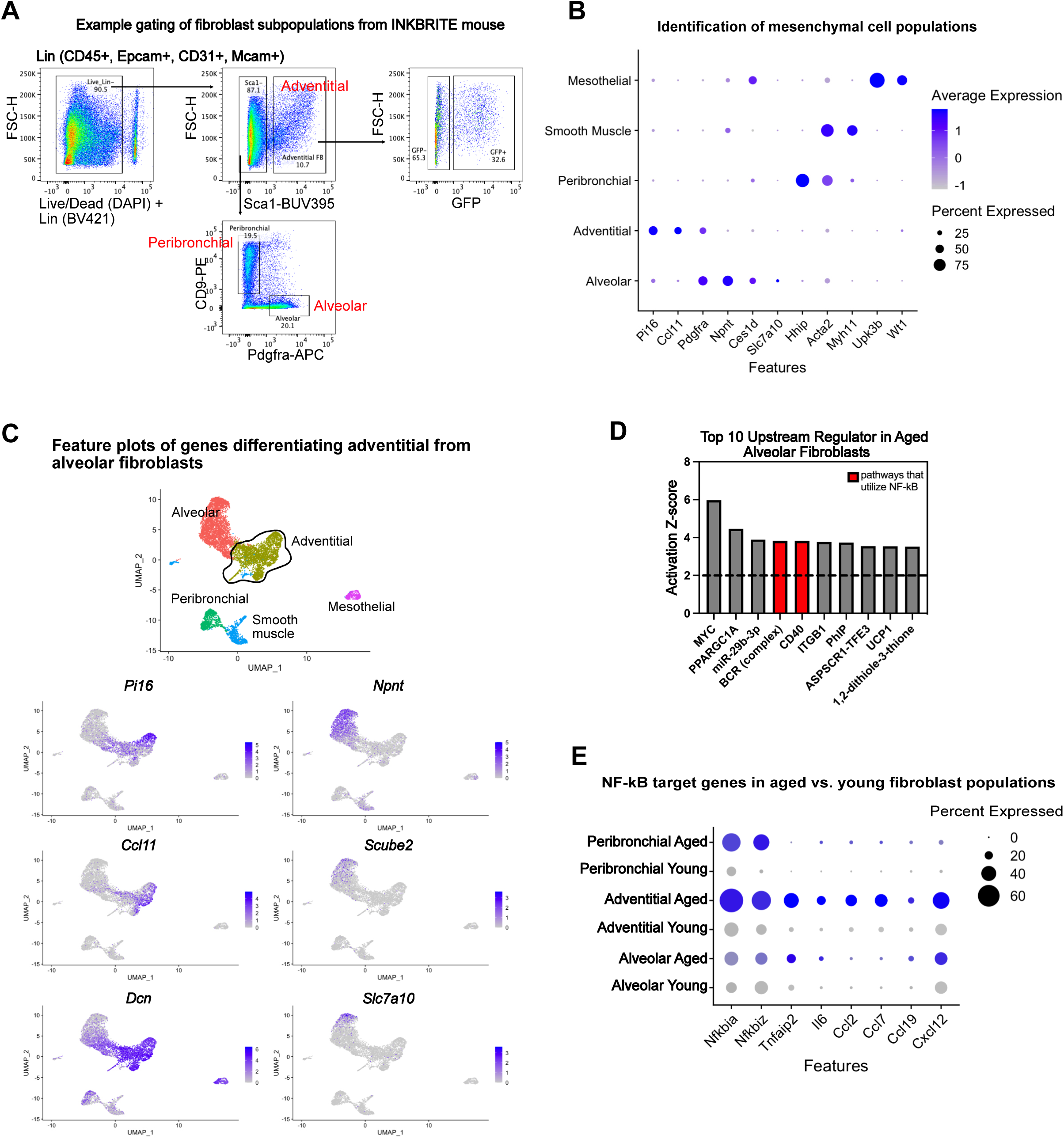
Evidence for enrichment of NF-kB pathway activation within aged adventitial fibroblasts. (A) Flow cytometry gating strategy for fibroblast subpopulations in INKBRITE (p16^INK4a–^GFP reporter) mice. (B) Dotplot of genes used to identify fibroblast and other mesenchymal subpopulations from scRNA sequencing of mesenchyme (CD45-CD31-Epcam-) from young (2 mo.) and aged (22 mo.) WT C57BL/6 mice. (C) Feature plots of genes used to differentiate adventitial and alveolar fibroblasts. (D) Qiagen Ingenuity Pathway Analysis (IPA) of lung fibroblast scRNA-seq comparing predicted upstream regulators within aged versus young alveolar fibroblasts. An activation score >2 predicts pathway activation. (E) Dot plot of NF-kB target genes by fibroblast type and age.

**Figure S3.**
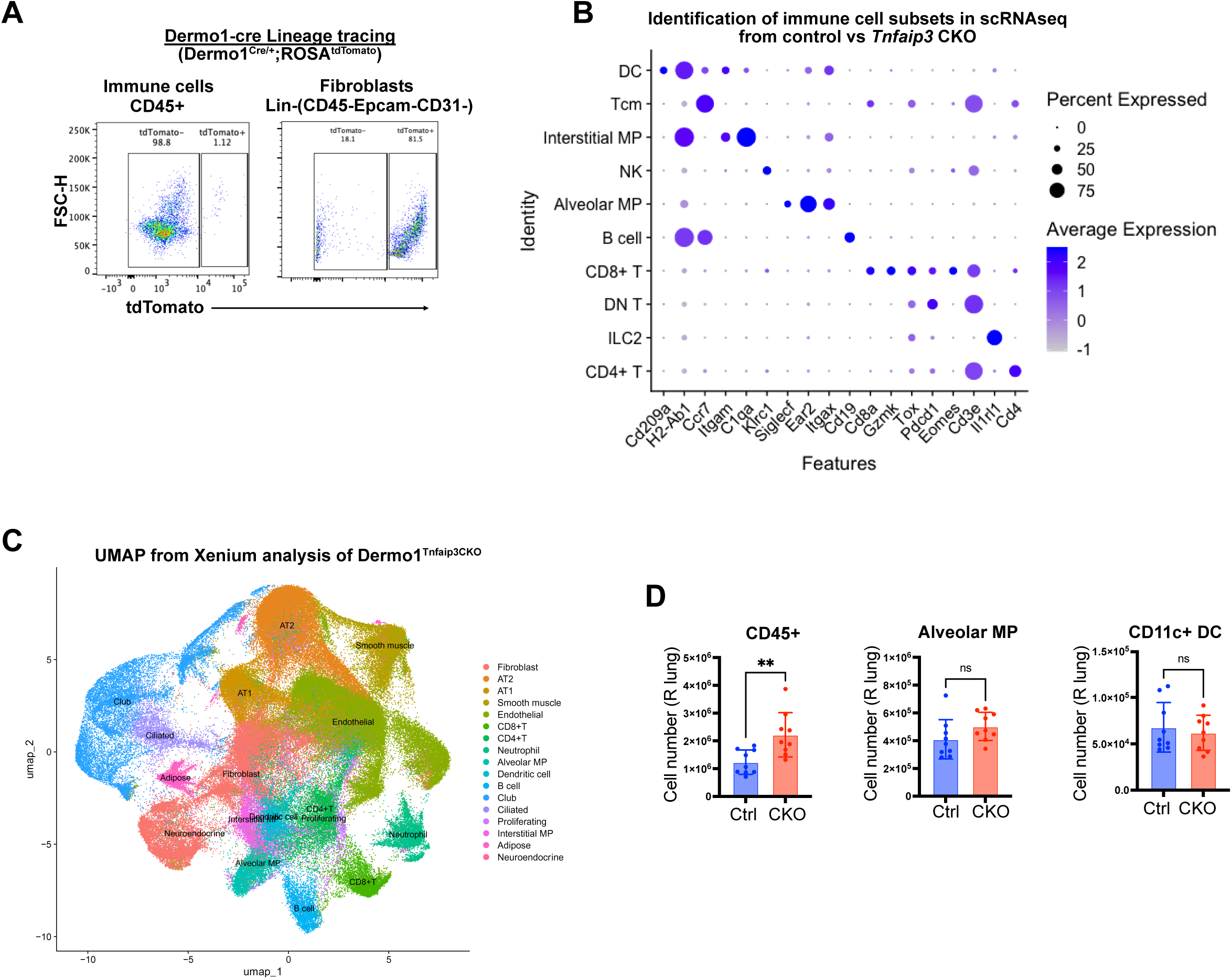
Fibroblast deletion of *Tnfaip3* in young mice leads to the expansion of lung resident CD8^+^ T cells. (A) Activation of lineage reporter (tdTomato) in fibroblasts compared to CD45^+^ immune cells in *Dermo1^cre/+^:R26R^tdTomato^* lungs. (B) Dot plot of key genes used to identify immune cell subsets. (C) Seurat clustering and UMAP from Xenium data of 3 month *Dermo1^Tnfaip^*^3^*^−CKO^* lung. (D) Lung resident immune cell numbers in 3-month-old *Tnfaip3f/f* (Ctrl) or *Dermo1^Tnfaip^*^3^*^−CKO^* (CKO) mice as determined by flow cytometry (n=9 mice per group). Statistical analyses done by 2-tailed t-test. ** p<0.01.

**Figure S4.**
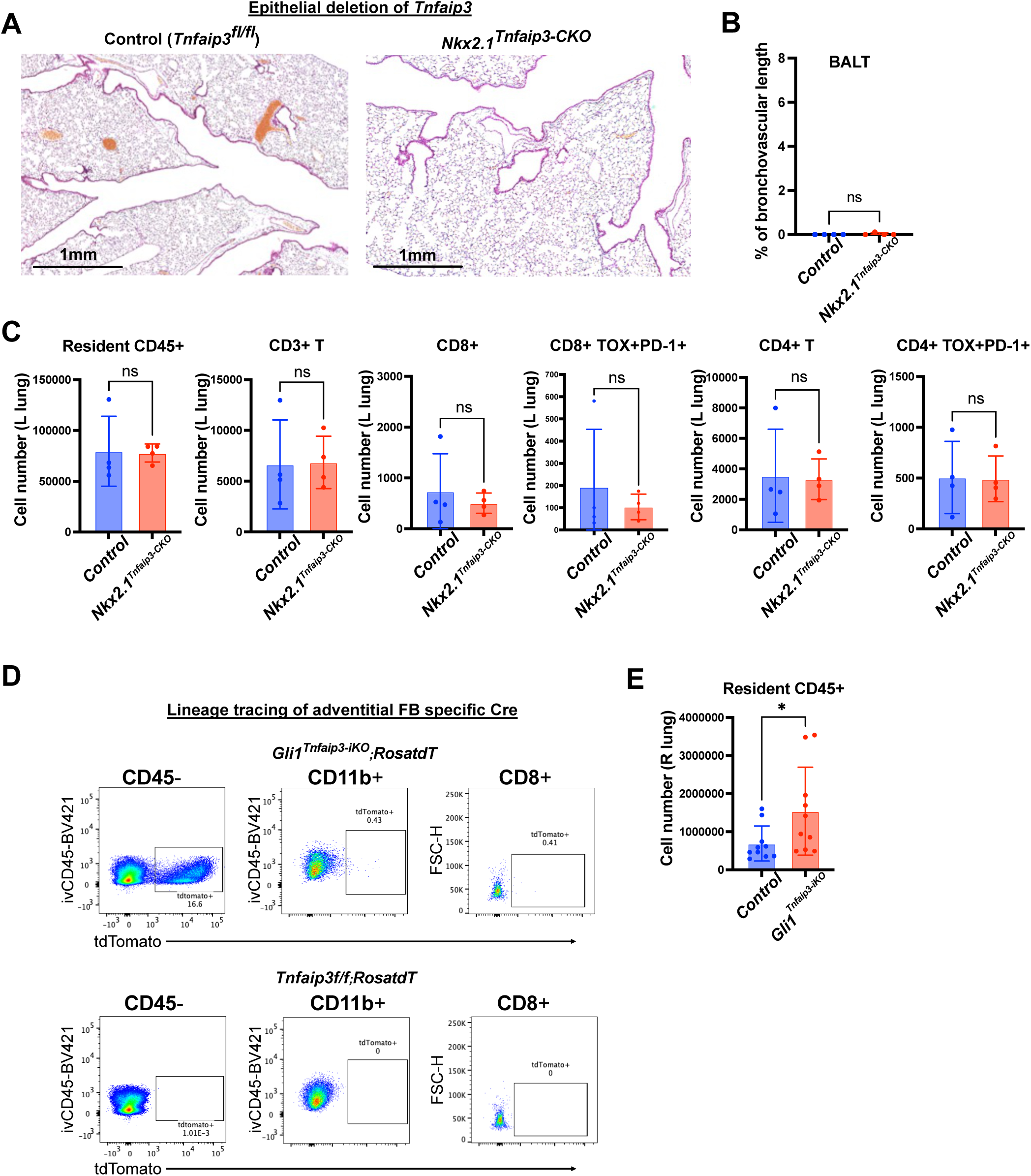
Deletion of *Tnfaip3* from adventitial fibroblasts but not epithelial cells contributes to inflammaging phenotype. (A-C) Analysis of 3 month 3 month old control (*Tnfaip3f/f*) and *Nkx2.1cre/+;Tnfaip3f/f (Nkx2.1^Tnfaip^*^3^*^−CKO^*) mice (n = 4 mice per group). (A) H&E staining of lung sections. (B) Quantification of BALT measured as percent of airways and vessels with adjacent lymphoid tissue. (C) Lung resident immune cell numbers determined by flow cytometry. (D) *Gli1*-creERT2 lineage tracing in indicated cell populations. (E) Lung resident CD45+ cell number in control vs *Gli1^Tnfaip^*^3^*^−iKO^* mice (n = 10 mice per group). Experimental design as described in Figure 3I. All statistical analyses done by 2-tailed t-test, except panel (E) which was analyzed by 1-tailed t-test. * p<0.05.

**Figure S5.**
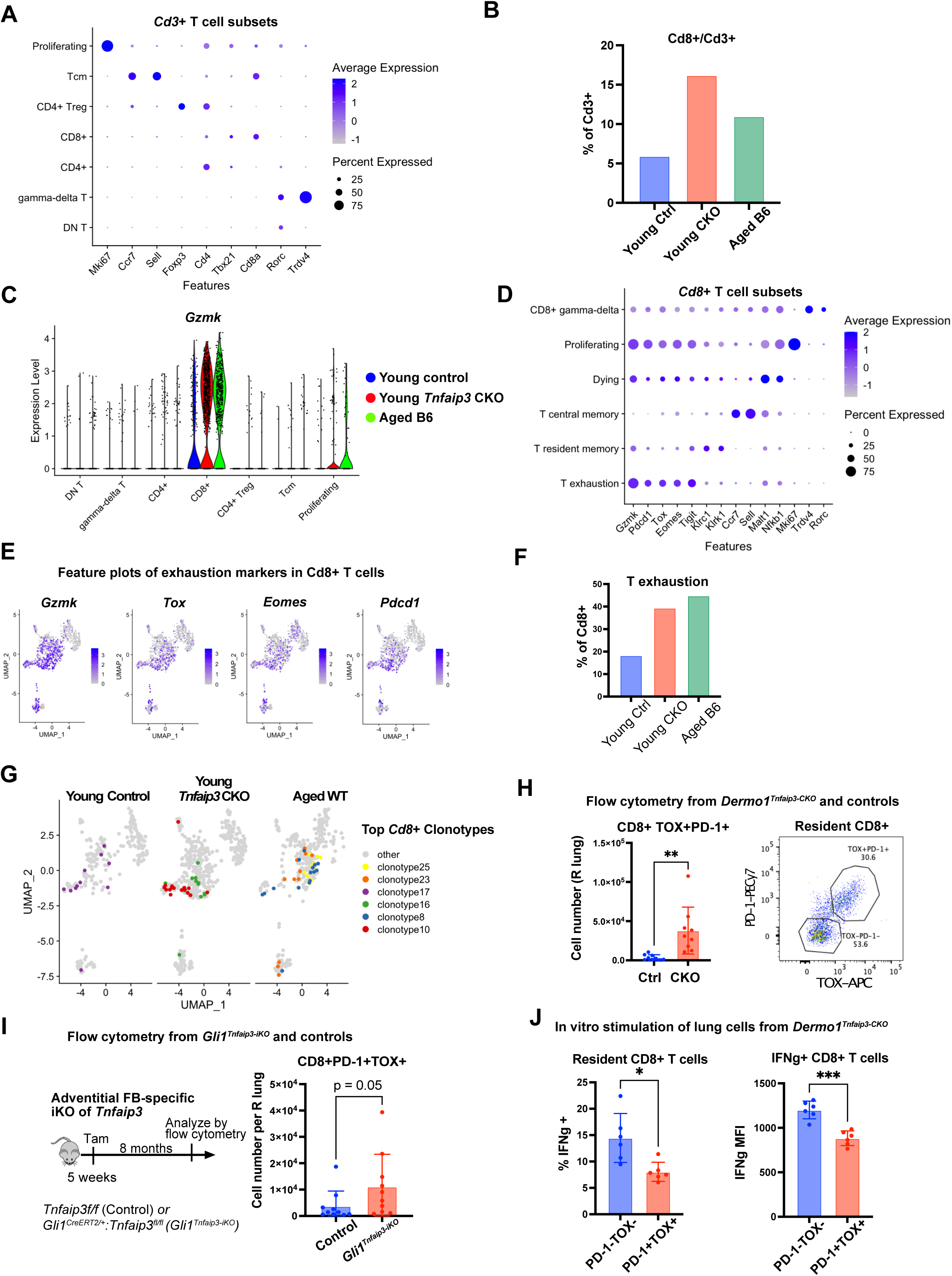
Fibroblast deletion of *Tnfaip3* in young mice phenocopies the expansion of an exhausted *Gzmk*+ CD8+ T cell population seen in normal aging. (A-G) scRNA-sequencing analysis of lung resident CD3+ T cells from young (3 month) control, young (3 month) *Tnfaip3* CKO and aged (23 month) B6 mice. (A) Dot plot depicting genes identifying T cell subsets. (B) *Cd8*+ T cells as a fraction of total *Cd3*+ cells. (C) Violin plot of *Gzmk* expression according to cell subset and sample. (D) Dot plot depicting genes identifying *Cd8*+ T cell subsets. (E) Feature plots of exhaustion genes within *Cd8*+ T cells. (F) Percent of *Cd8*+ T cells within exhaustion cluster. (G) TCR clonotypes with 10 or more cells sequenced per clonotype superimposed on *Cd8*+ UMAP. (H) CD8+TOX+PD-1+ lung resident T cells in young (3 mo.) control and *Dermo1^Tnfaip^*^3^*^−CKO^* mice, along with example gating (n = 9 mice per group). (I) Experimental schematic for adventitial fibroblast specific inducible knockout of *Tnfaip3*, along with number of lung resident CD8+TOX+PD-1+ cells at time of analysis (n = 10 mice per group). (J) Flow cytometric analysis of lung resident CD8+ T cells after 3 hours of in vitro stimulation with PMA, ionomycin and brefeldin A (n = 6 mice). Graphs depict percent of CD8+ T cells that are IFNg positive, as well as MFI of IFNg+ cells by TOX and PD-1 status. All statistical analyses are 2-tailed t-test, except for (I), which is a 1-tailed t-test. * p<0.05, ** p<0.01, *** p<0.001.

**Figure S6.**
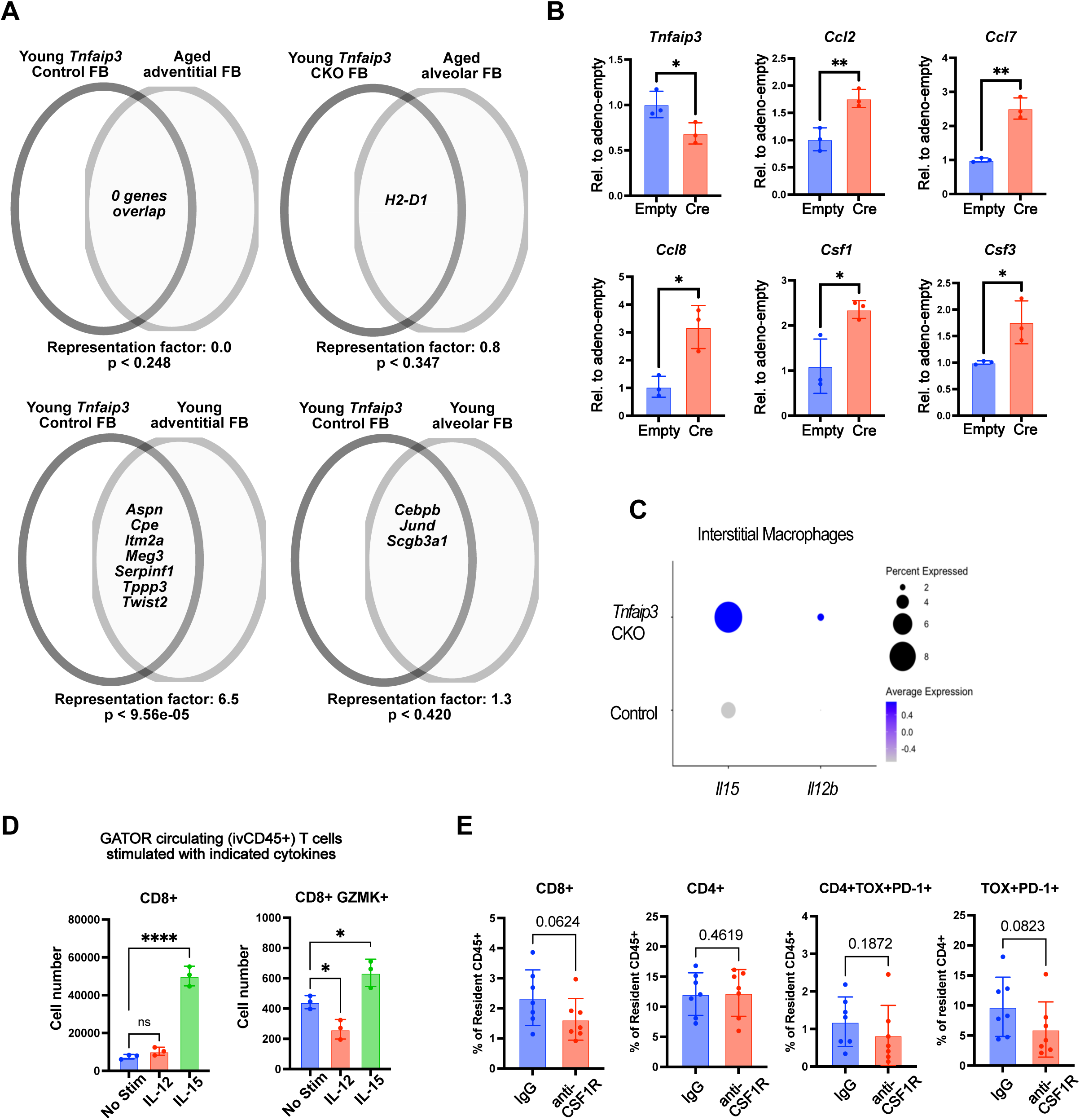
Fibroblast deletion of *Tnfaip3* promotes monocyte recruitment which in turn drives T cell exhaustion through the secretion of cytokines. (A) Venn diagram demonstrating overlap of genes upregulated in both fibroblast populations relative to their counterparts (e.g. aged alveolar FB vs. young alveolar FB, Tnfaip3 CKO FB vs. control FB). A representation factor > 1 indicates more overlap than expected between two independent groups. (B) qPCR expression analysis of *Tnfaip3* and other NF-kB target genes in *Tnfaip3f/f* lung fibroblasts infected with either an empty or cre-expressing adenovirus (Cre) (n = 3 wells per condition). (C) Dot plot showing expression of indicated genes in interstitial macrophage subset from scRNA sequencing of 2 month *Tnfaip3f/f* (Control) and *Dermo1^Tnfaip^*^3^*^−CKO^* mice (D) Day 5 analysis of circulating T cells from GATOR mice incubated with indicated cytokines. Cell numbers determined by flow cytometry. (E) Indicated immune populations within *Dermo1^Tnfaip^*^3^*^−CKO^* mice as a percentage of resident immune and CD4+ T cells after 2 months of antibody treatment. Full experimental schematic described in Figure 5K. Panel (B) analyzed by 2-tailed t-test, panel (D) by 1-way ANOVA, and panel (E) by 1-tailed t-test. * p<0.05, ** p<0.01, **** p<0.0001.

**Figure S7.**
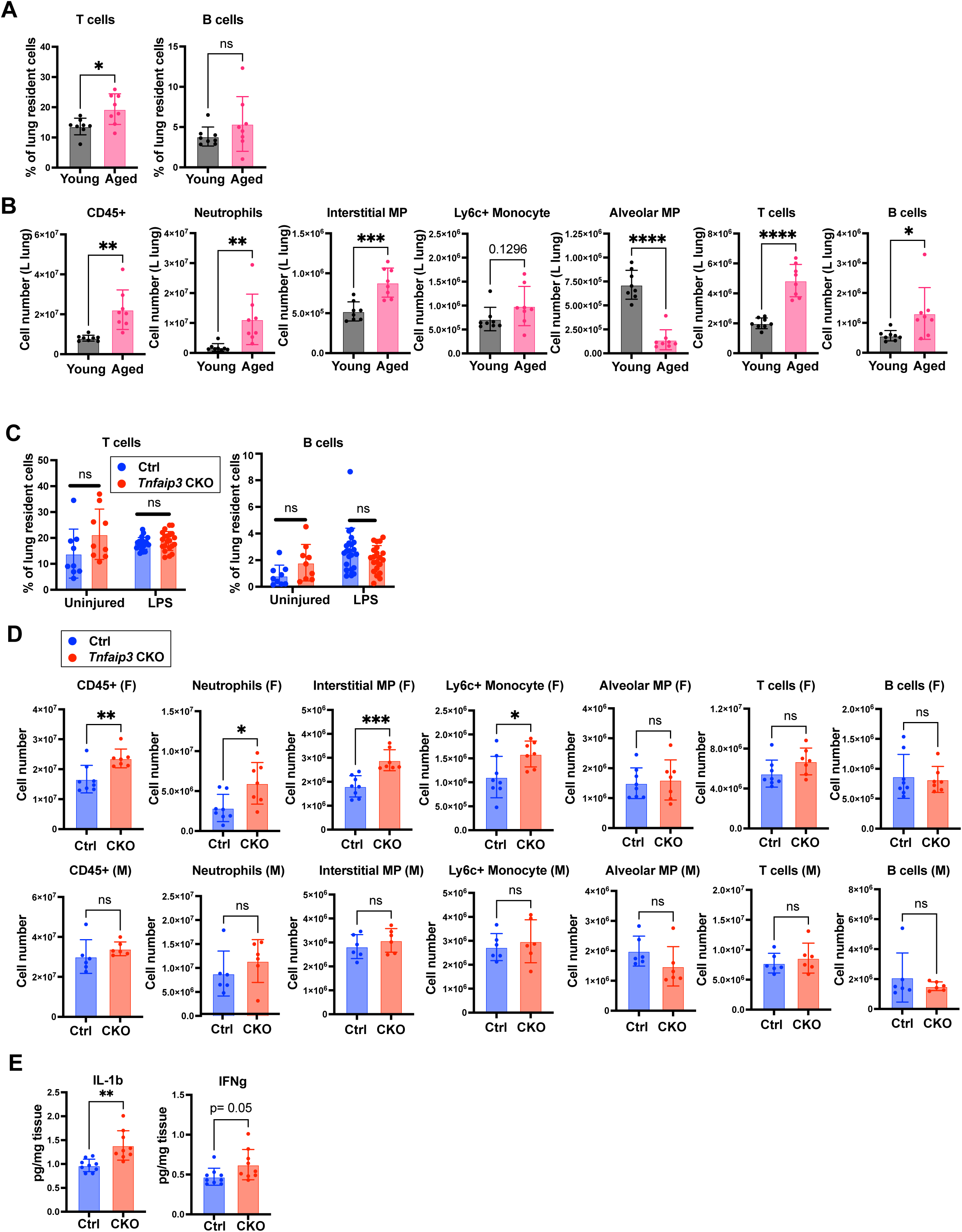
Fibroblast deletion of *Tnfaip3* phenocopies the exaggerated inflammatory response observed in aged animals in a model of ARDS. (A-B) Flow cytometric analysis of lung resident immune cells day 4 post-LPS in aged and young B6 mice. (A) Lymphocyte populations as a fraction of all lung cells and (B) cell number of indicated lung resident immune cells (n = 8 female mice per group). (C) Lymphocyte populations as a percentage of total lung resident cells in control or *Tnfaip3* CKO mice day 4 post-LPS (n = 21-22 mice per group). Percentages of lymphocyte populations in homeostasis from a similarly age and sex-matched cohort included for comparison (n = 9 mice per group). (D) Cell number of indicated lung resident immune cells in control or *Tnfaip3* CKO mice on day 4 post-LPS, separated by sex (control male n = 6, female n = 8, CKO male n= 6, female n = 7). (E) Indicated protein concentrations from lung homogenates of control and *Tnfaip3* CKO mice on day 4 post-LPS as determined by Luminex assay (n = 9 mice per group). Statistical analyses all done by 2-tailed t-test. * p<0.05, ** p<0.01, *** p<0.001, **** p<0.0001.

**Figure S8.**
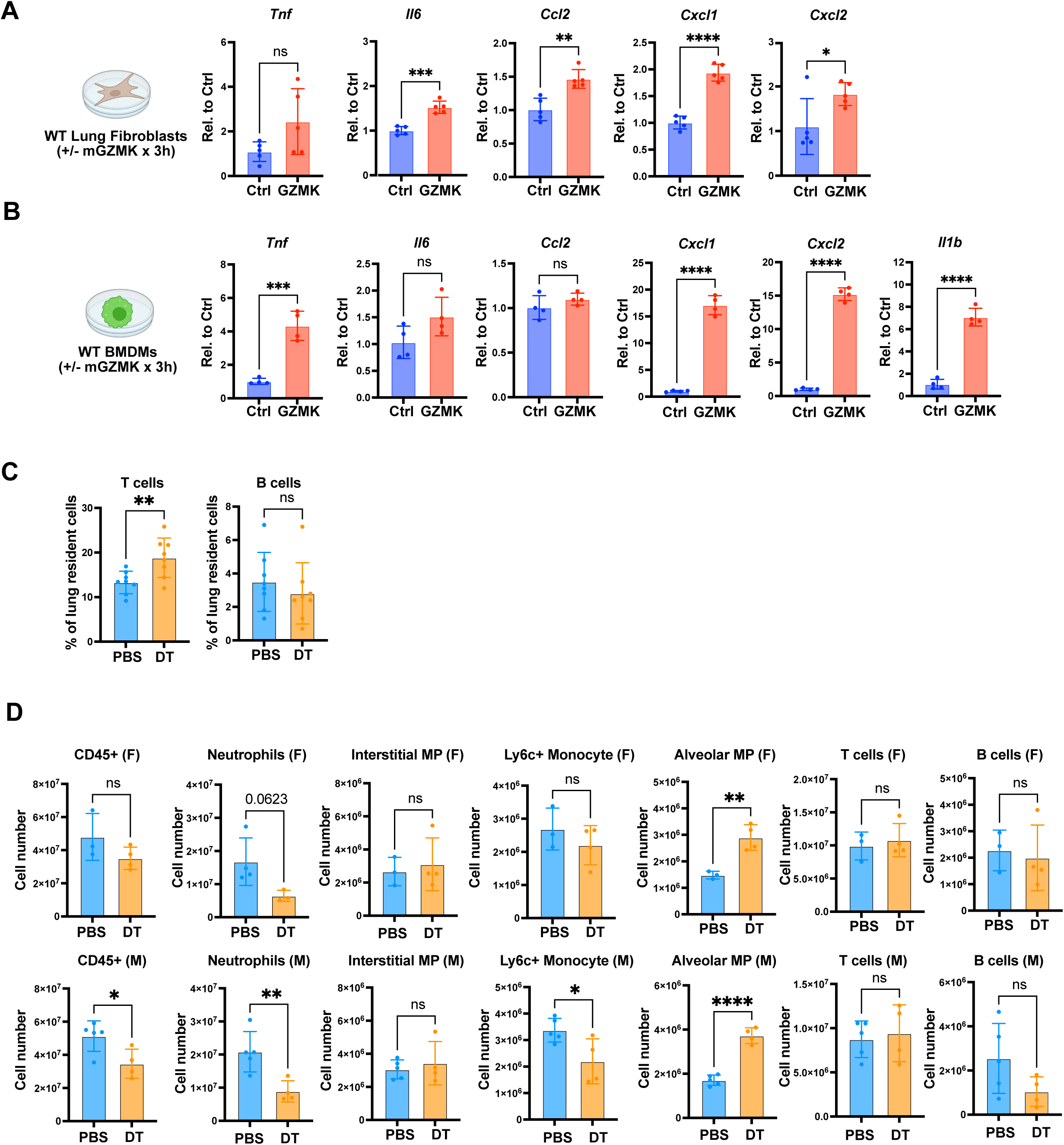
Granzyme K potentiates lung inflammation. (A) qPCR expression analysis of indicated genes in wildtype lung fibroblasts cultured with or without recombinant mouse GZMK at a concentration of 100ng/mL for 3 hours (n = 5 wells per condition). (B) qPCR expression analysis of indicated genes in wildtype bone marrow derived macrophages cultured with or without recombinant mouse GZMK at a concentration of 100ng/mL for 3 hours (n = 4 wells per condition). (C-D) Flow cytometric analysis of *Dermo1^Tnfaip^*^3^*^−CKO^;*GATOR mice day 4 post intranasal LPS that had either received PBS or diptheria toxin (DT) prior to LPS, as described in Figure 6I. (C) Lymphocyte populations as a fraction of all lung cells and (D) cell number of indicated lung resident immune cells, separated by sex (PBS male n = 5, female n = 3, DT male n = 4, female n = 4). All statistical analyses performed with 2-tailed t-test. * p<0.05, ** p<0.01, *** p<0.001, **** p<0.0001.

**Figure S9.**
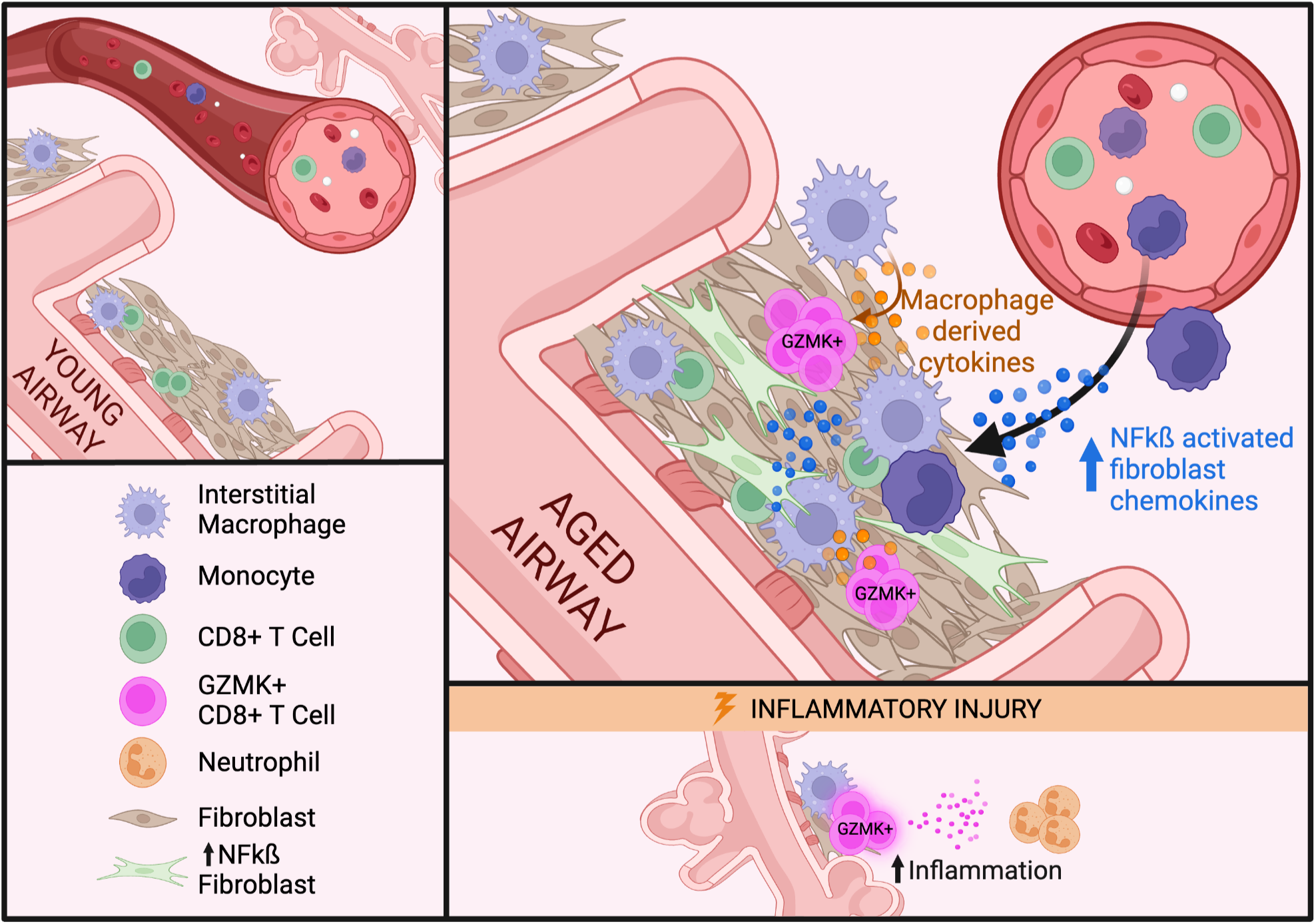
Model of fibroblast orchestration of inflammaging via NF-kB activation. (LEFT, TOP) In youth, lung adventitial fibroblast NF-kB activation is low during homeostasis. (RIGHT, TOP) With age, there are homeostatic elevations in NF-kB activation within adventitial fibroblasts, increasing the expression of monocyte chemotactic factors and supporting the expansion of the lung resident interstitial macrophage population. The expanded interstitial macrophage population in turn drives the proliferation and ultimately exhaustion of resident CD8+ T cells through the secretion of cytokines, creating a large reservoir of GZMK+ CD8+ T cells within BALTs of aged lungs. (RIGHT, BOTTOM) In the setting of an inflammatory stimulus these GZMK+ CD8+ T cells augment tissue inflammation, contributing to the increased risk of older individuals to ARDS and potentially other inflammatory lung diseases.

## MATERIAL AND METHODS

### Animal Studies

All animals were housed and treated in accordance with the Institutional Animal Care and Use Committee (IACUC) protocol approved at the University of California, San Francisco Laboratory Animal Resource Center (LARC). All animals used in this study were maintained in the C57BL/6 background. The GATOR mice were generated by knocking in a cassette with homology arms flanking a T2A-CreERT2-T2A-tdTomato-P2A-DTR insert to replace the TGA stop codon in exon 5 of the mouse *Gzmk* locus. Genotype was confirmed in F1 animals by PCR screening of the 5’ and 3’ arm of the inserted cassette into the endogenous locus. Generation and genotyping of the *Gli1^cre/ERT^*^2^, *R26R^tdTomato^*, *Nkx2.1^cre^, Dermo1^cre^*mouse lines were performed as previously described by The Jackson Laboratory. The *Tnfaip3^flox/flox^* mouse line was generated in the Ma lab and the INKBRITE reporter was generated in the Peng lab. For lineage tracing studies, tamoxifen was dissolved in corn oil and administered intraperitoneally (IP) at 200 mg/kg body weight per day for 5 consecutive days. Lipopolysaccharide (LPS) (Sigma, cat# L2880-25MG) was delivered intranasally once at 1mg/kg body weight in 50µl of sterile 1X PBS. For depleting *Gzmk*^+^ cells *in vivo*, diphtheria toxin (Sigma, D0564) or vehicle (sterile PBS) was administered intraperitoneally at 50 ng/g of mouse per day x 5 days. For interstitial macrophage depletion either 50ug of anti-CSF1R (BioXCell BP0213) or rat IgG2a control (BioXCell BP0089) was administered intranasally in 50ul sterile PBS twice weekly for 8 weeks. For all intranasal administrations mice were anesthetized with isofluorane.

### Tissue Dissociation and Flow Cytometry

For analysis of lung fibroblasts, dissected mouse lungs were tracheally insufflated with a digestion cocktail of Collagenase Type I (225 U/mL, Thermo Fisher), Dispase (15 U/mL, Thermo Fisher), 1x Pen/Strep (Thermo Fisher, 15070063) and Dnase (50 U/mL, Sigma), and removed from the chest. The lung was then incubated in an additional 5 mLs of digestion buffer for 45 mins at 37 °C in a shaker at 200 rpm. The mixture was then washed with FACS buffer (2% FBS, 1x Pen/Strep in DMEM w/o phenol red) and passed through a 70 µm cell strainer prior to centrifugation at 550g x 5 minutes. The cell pellet was then resuspended in 1x RBC lysis buffer for 2 minutes prior to dilution with FACS buffer and passage through a 40 µm cell strainer and repeat centrifugation. Cells were then blocked with mouse FcR blocking reagent (Miltenyi Biotec 130-092-575) for 15 minutes at 4°C. To enrich for fibroblasts prior to analysis or sorting, cells were incubated with biotin-conjugated antibodies as follows: CD146 (MCAM)-Biotin (Biolegend 134716, clone ME-9F1), CD45-Biotin (Biolegend 103104, clone 30-F11), CD326 (EpCAM)-Biotin (Biolegend 118204, clone G8.8), and CD31 (PECAM-1) (Invitrogen, 13-0311-82, clone 390), with subsequent magnetic bead depletion of these labeled cells using the EasySep Mouse Streptavidin Rapidspheres Isolation Kit (StemCell Technologies, 19860). Cells that were not depleted were subsequently incubated with the appropriate antibodies in FACS buffer for 30 min at 4°C and washed with FACS buffer. Antibodies used for the analysis of fibroblasts are as follows: Live/Dead (DAPI, Fisher Scientific D1306) and Dump channel included CD326 (EpCAM)-BV421 (BD Biosciences 563214, clone G8.8), CD31-ef450 (Invitrogen 48031182, clone 390), Streptavidin-BV421 (BD Biosciences 563259), and CD45 BV-421 (BD Biosciences, clone 30-F11). Fibroblast markers included Sca1-BUV395 (BD Biosciences 563990, clone D7), PDGFRa-APC (Invitrogen 17140181, clone APA5) and CD9-PE (Millipore Sigma SAB4700, clone EM-04).

For the analysis of lung immune cells, 3 min before euthanizing each mouse 3 μg of anti-CD45-BV421 antibody (BD Biosciences, clone 30-F11) diluted in 150 μl of 1X sterile DPBS was intravenously injected through retro-orbital injections of isofluorane anesthetized mice. After euthanasia, dissected mouse lungs were tracheally insufflated with a digestion cocktail of Liberase TM (40 µg/mL, Sigma), DNase (50 U/ml, Sigma) and 1x Pen/Strep (Thermo Fisher, 15070063) in HBSS (+Ca+Mg). The lung was then incubated in an additional 5 mLs of digestion buffer for 45 mins at 37 °C on a shaker at 200 rpm. The mixture was then washed with FACS buffer (2% FBS in DMEM w/o phenol red) and passed through a 70 µm cell strainer prior to centrifugation at 550g x 5 minutes. The cell pellet was then resuspended in 1x RBC lysis buffer for 2 minutes prior to dilution with FACS buffer and passage through a 40 µm cell strainer and repeat centrifugation.

Prior to staining cells counts were obtained using a NucleoCounter (Chemometic). For experiments that tested cytokine production, 2-3×10^6^ cells were plated in 96-well U-bottom plate and stimulated ex vivo with 1X Stimulation Cocktail (Cytek, TNB-4975-UL100) in complete media (Advanced RPMI1640, 10% FBS, 1x GlutaMAX, 100 U/mL penicillin/streptomycin (1x P/S) and 50 µM β-mercaptoethanol) for 3 hours at 37°C prior to collection.

Cells were then blocked with mouse FcR blocking reagent (Miltenyi Biotec 130-092-575) for 15 minutes at 4°C and incubated with the appropriate extracellular antibodies in FACS buffer for 30 min at 4°C and washed with FACS buffer. The following extracellular antibodies were used for staining: CD3-FITC (Invitrogen 11003282, clone 17A2), CD45-BUV395 (BD Biosciences 564279, clone 30-F11), CD19-PE/Dazzle594 (Biolegend 115554, clone 6D5), CD8-PE (Biolegend 100798, clone 53-6.7), CD8-BV786 (BD Biosciences 563332, clone 53-6.7), PD-1-PE-Cy7 (Biolegend 109110, clone RMP1-30), TCRgd-PerCP-Cy5.5 (Biolegend 118117, clone GL3), SiglecF-BV786 (BD Biosciences 740956, clone E50-2440), CD11c-PE-Cy7 (Biolegend 117318, clone E50-2440), Ly6g-APC-Cy7 (Biolegend 127624, clone 1A8), CD11b-BV711 (Biolegend 101241, clone M1/70), CD3-PE/Dazzle594 (Biolegend 100246, clone 17A2), I-A/I-E (MHCII)-BUV737 (BD Biosciences 748845), clone M5114), CD64-AF647 (BD Biosciences 558539, clone X54-5/7.1), Ly6c-AF488 (Invitrogen 53593282, clone HK1.4), and CD11b-BV605 (BD Biosciences 563015, clone M1/70). For experiments involving intra-cellular staining, cells were fixed and permeabilized using the Foxp3 Staining Buffer Set (Thermo Fisher, 00-5523-00) after extracellular staining was performed. Antibodies utilized for intracellular staining include IFNg-APC (Invitrogen 17-7311-81, clone XMG1.2), TOX-ef660 (Invitrogen 50-6502-82, clone TXRX10) or TOX-PE (Invitrogen 12-6502-82, clone TXRX10).

Cell analysis was performed on a 5-laser BD Fortessa. Cell sorting was performed on a 5-laser BD FACSAria Fusion or BD FACSAria II. All instruments used FACSDiva Software. FlowJo 10.8.1 software was used for analysis.

### Single-cell RNA sequencing and TCR sequencing analysis

For sample submission of lung resident immune (ivCD45-,CD45+) cells from young control and young *Dermo1^Tnfaip^*^3^*^−CKO^* animals, cells from 3 independent mice were combined per genotype. For sample submission of young and aged lung fibroblasts (CD31-,EPCAM-,CD45-), cells from a single aged and young C57BL/6 mouse were submitted. For sample submission of lung resident T cells (ivCD45-,CD45+,CD3+) from young control, young *Dermo1^Tnfaip^*^3^*^−CKO^*, and aged C57BL/6 animals, cells from 3 independent mice were combined per sample.

Single cell sequencing was performed on a 10X Chromium instrument (10X Genomics) at the Institute of Human Genetics (UCSF, San Francisco, CA). Briefly, live mouse lung cells were sorted and resuspended at a concentration of 1,000 cells/µl in 50 µl PBS with 0.04% BSA prior to being loaded onto a single lane of the Chromium Controller to produce gel beads-in emulsion (GEMs). GEMs underwent reverse transcription for RNA barcoding and cDNA amplification. For lung fibroblast and lung resident CD45+ immune cell sequencing (those that did not use TCR sequencing), the library was prepped with the Chromium Single Cell 3’ Reagent Version 3 kit. For single cell sequencing of lung resident T cells from young control (*Tnfaip3f/f*), young Dermo1^Tnfaip3f/f^ , and aged C57BL/6 control mice, the Chromium Next GEM Single Cell 5’ Reagent Kits v2 were used to prepare the library for gene expression, TCR, and cell surface hashtag sequencing. Biolegend hashtag oligos were added to their respective samples (Young control: TotalSeq-C0301, Young Dermo1^Tnfaip3f/f^: TotalSeqC0302, and Aged C57BL/6: TotalSeqC0303) during cell surface staining, prior to sorting. Paired-end sequencing was performed on an Illumina NovaSeq6000. Raw Fastq files were aligned with Cellranger. Demultiplexing was performed in Seurat using HTODemux. We used the Seurat R package along with a gene-barcode matrix provided by CellRanger for downstream analysis. Following the standard workflow of Seurat, we generated Seurat objects after using ScaleData, RunPCA, RunUMAP.

### Bulk RNA Sequencing and Analysis

Bulk RNAseq sequencing was done using Sanger/Illumina 1.9 with a total of 3 biological replicates per condition. Quality control of reads was conducted by using FastQC (Babraham Bioinformatics). Ligation adaptors were removed using Cutadapt and Sickle. Sequencing reads were aligned using HISAT and assembled with Stringtie software to the reference genome *Mus musculus*, UCSC version mm10. All gene counts of the biological replicates were concatenated while running DEseq2 for differential gene expression (DGE). To assess the enrichment for genes upregulated in both the lung fibroblasts of *Dermo1^Tnfaip^*^3^*^−CKO^* samples (compared to controls) and upregulated in aged adventitial fibroblasts (compared to young adventitial fibroblasts), we calculated the statistical significance of the overlap between the 2 genes sets utilizing the webtool on nemates.org, which calculates the representation factor along with the p value of overlap as a measure of the significance of enrichment utilizing hypergeometric probability testing.

### Xenium sample preparation

Xenium analysis was performed on one lung section from a 24-month old female INKBRITE mouse, and one lung section from a 3-month female *Dermo1^Tnfaip3CKO^*mouse. To prepare tissue for analysis, 5 μm FFPE tissue were placed onto a Xenium slide, followed by deparaffinization and permeabilization to make the mRNA detectable. The Xenium platform from 10X Genomics was used according to the manufacturer’s recommendations and as previously reported^58^. 379 genes from the Xenium Mouse Tissue Atlassing panel in addition to a 50 gene custom panel were used for analysis.

Custom panel gene list is as follows: Itgam, Itgax, C1qa, Il15, Il12b, Il18, Cx3cr1, Ccr2, Ly6c1, Siglecf, Cd4, Tox, Pdcd1, Gzmk, Ifng, Gzmb, Eomes, Mki67, Tcf7, Ccl2, Ccl5, Ccl7, Pdgfra, Il6, Hhip, Gli1, Tnf, Cd19, Krt5, Shh, Trp63, Muc5ac, Cdkn2a, Il7, Apoe, Acta2, Postn, Cthrc1, Cxcl12, Ly6a, Tigit, Hnf4a, Acadl, Fads3, Ffar4, Lpin2, Fasn, Far1, tdTomato, and eGFP.

### Spatial cluster generation and mapping from Xenium

We employed the Seurat vignette (https://satijalab.org/seurat/reference/readxenium) to load and analyze the Xenium data with Seurat version 5. For normalization, we applied SCTransform method, followed by standard dimensionality reduction and clustering. The clustering results were visualized in UMAP space. Subsequently, we annotated each cluster according to their gene expression. The annotated clusters were imported into the Xenium explorer to map their spatial locations.

### Cell neighborhood analysis and BALT composition analysis in spatial transcriptomic data

The spatial coordinates of cell centroids (defined by the Xenium instrument) were organized into a K-D tree, and each cell’s fixed-radius neighbors were identified using the dbscan package in R. For BALT composition analysis cells from 4 BALTs per section were combined for analysis.

### Tissue preparation and immunohistochemistry

For paraffin embedded mouse lungs, mouse right ventricles were perfused with 1 ml PBS and the lungs were inflated with 4% PFA, and then fixed in 4% PFA overnight at 4°C. After fixation, the lungs were washed by cold PBS X 4 times over 2 hrs at 4°C and dehydrated in a series of increasing ethanol concentration washes (30%, 50%, 70%, 95% and 100%), each a minimum of 2 hrs. The dehydrated lungs were incubated with Xylene for 1 hr at RT and with paraffin at 65°C for 90 min X 2 times, and then embedded in paraffin and sectioned. For staining lung sections were deparaffinized in xylene followed by stepwise rehydration from 100% ethanol to PBS. Antigen retrieval was performed with 1x Diva Decloaker (Biocare Medical, DV2004) according to protocol. Sections were then permeabilized with PBS with 0.1% Tween20 (PBST) and blocked with 3% donkey serum in PBST for 1 hour at RT prior to overnight incubation at 4°C with primary antibodies in PBST with 1% donkey serum. The following primary antibodies were used: tdTomato (Sicgen, 1:200, AB8181-200) and CD3 (1:300, Abcam, ab16669). Following overnight incubation with primary antibody, sections were washed in PBST x 3 and incubated with fluorescently-conjugated secondary antibodies in PBST for 1 hour at RT. Secondary antibodies used include Donkey anti-Rabbit IgG Alexa Fluor 488 (Thermo-Fisher, A-21206) and Donkey anti-Goat IgG Alexa Fluor 555 (Abcam, ab150130). Sections were again washed with PBST, nuclei were stained with DAPI, and slides were treated with Trueblack Autofluor Quencher (Biotium, 23007) prior to a final wash with PBS and mounting with FluorSave Aqueous Mounting Medium (VWR, EM345789-20ML).

For O.C.T (Scigen 4586) embedded lungs, after mouse right ventricles were perfused with PBS and insufflated with 4% PFA, lungs were fixed in 4% PFA for 2 hrs at 4°C and then washed four times over 2 hours in PBS. Lungs were then placed in 30% sucrose in PBS and incubated at 4°C overnight, followed by overnight incubation at 4°C in a 1:1 mixture of 30% sucrose in PBS and O.C.T. Lungs were then incubated in 100% O.C.T for 3 hours prior to embedding in O.C.T. and cyrosectioning. All incubation steps were done with slow rocking of samples. For O.C.T embedded sections, rehydration and antigen retrieval steps were skipped. Otherwise, sample processing was as described under paraffin sectioning. Primary antibodies used for cryosectioned lungs include CD3 (1:300, Abcam, ab16669) and CD11b (1:200, Biolegend 101248). Secondary antibodies include Donkey anti-rabbit Alexa Fluor 647 (Thermo Fisher, A78967) and Donkey anti-rat Alexa Fluor 488 (Thermo Fisher, A48269). Signal for Gli1 lineage tracing from *Gli1^Tnfaip^*^3^*^−iKO;RosatdT^* mice was endogenous tdTomato signal.

### Histological analysis of BALT and quantification of alveolar damage

For morphological analysis of tissue, paraffin embedded sections were deparaffinized as previously described and stained according to standard protocols with hematoxylin and eosin prior to dehydration and coverslip mounting with Cytoseal (VWR, 48212-154).

For BALT quantification, lung sections were imaged on a Molecular Devices Image Xpress PICO Automated Cell Imaging machine at 4x magnification. To quantify BALT, ImageJ2 (Version 2.14.0) was used to calculate the total length of airways and vessels in a full lung section containing large airways. The length of the airways and vessels with adjacent lymphoid tissue was then measured in a similar manner and BALT was quantified as percent of airways and vessels with adjacent lymphoid tissue.

For assessment of LPS-induced alveolar damage, histological analysis was performed by a board-certified comparative pathologist blinded to experimental groups. H&E images were analyzed using Olympus cellSens software. The severity and extension of alveolar damage was assessed using a semiquantitative grading system described by Matute-Bello et al. (2010) that includes criteria like polymorphonuclear cell infiltration, alveolar and interstitial, presence of hyaline membranes, fibrin strands and thickening of alveolar septa. 20 random histological fields (40x magnification) of lung were analyzed according to these criteria per mouse and a semi-quantitative score was generated, as depicted in Figure 6F.

### Cellular p65 staining and Intensity Quantification

Primary adventitial fibroblasts were sorted from aged (>24 months) and young (2 month) male C57BL/6 mice according to gating strategy described in Figure S2. Cells were plated on a sterile laminin coated coverslip (Fisher Scientific, NC0597572) and cultured for 3 days in standard media (DMEM, 10% FBS, 1%Pen/Strep) to allow for cellular adherence. Cells were then fixed in 4% PFA for 15 minutes at RT. After fixation, cells were rinsed with PBST and permeabilized in 0.2% Triton X-100 in PBS for 15 minutes at RT prior to again washing with PBST. Cells were blocked for 1 hour in PBST with 3% donkey serum and Donkey anti-mouse IgG (Jackson ImmunoReserach, 715-007-003) at 1:20 dilution. Cells were stained overnight at 4°C with anti-p65/RELA (1:500, Santa Cruz Biotechnology, sc-8008) in PBST with 1% donkey serum. The following day cells were washed with PBST and stained with donkey anti-mouse Alexa Fluor 555 (Invitrogen, A-31570) secondary antibody prior to staining with DAPI and mounting with FluorSave Aqueous Mounting Medium (VWR, EM345789-20ML). Images were acquired using a fluorescence microscope at 10x magnification. Image processing was conducted in FIJI, where raw .zvi files were separated into grayscale images for the DAPI and p65 channels. Nuclear location and morphology were defined by DAPI staining, and thresholding segmented these regions by excluding dim signals (background noise), bright signals (artifacts), and cytoplasmic areas. The binary watershed algorithm resolved adjoining nuclei without manual intervention. Non-cellular particles (<7 pixels²) and dying fibroblasts (p65 oversaturation and contracted morphology) were deselected. The live-cell nuclear regions were added to the ROI Manager for p65 intensity analysis. For optimal signal contrast, the p65 channel brightness was adjusted consistently across all images. The selected DAPI regions were superimposed onto the p65 channel along with selection of a neighboring cytoplasmic regions with an average non-nuclear p65 intensity. The mean intensity of selected regions was measured and the ratio of nuclear to cytoplasmic p65 intensity was calculated, where values >1 indicating higher nuclear expression. More than 150 cells per sample were analyzed.

### Thick section imaging and cell distance quantification

For detection of vascular-associated leukocytes, mice were injected with 3 μg of BV421-conjugated CD45 antibody by tail vein injection 3 mins prior to euthanasia. After euthanasia and dissection, the right ventricle was gently perfused with PBS prior to tracheal insufflation with 4% PFA and overnight fixation at 4°C. After washing with 1X DPBS, lung lobes were placed in 30% sucrose for 3 days then frozen in O.C.T. (Thermo Scientific). Coronal sections of 200 mm were prepared using a cryostat (Leica). Samples were washed extensively with 1xPBS and incubated in Ce3D staining/blocking buffer (DPBS/0.3%Triton X-100/1% BSA/1% horse serum) for 1 day at room temperature. After, samples were incubated with unconjugated primary antibodies diluted in in Ce3D staining/blocking buffer (DPBS/0.3%Triton X-100/1% BSA/1% horse serum) at room temperature 1-2 days. Primary antibodies used for this study include CD8-AF647 (Biolegend 100727, 1:100), Goat anti-RFP (Rockland, 600-901-379, 1:200), Rabbit anti-Iba1 (Wako 019-19741, 1:400), and aSMA AF594 (Abcam ab202368, 1:300). Next, samples were washed in Ce3D washing buffer (DPBS/0.2% Tween-20/thioglycerol) for 30 min, 3–4 times, then incubated with secondary antibodies and conjugated primary antibodies diluted in Ce3D staining/blocking buffer (DPBS/0.3%Triton X-100/1% BSA/1% horse serum) at room temperature 1-2 days. Secondary antibodies used include Donkey anti-Rabbit AF488 (Life Technologies A21206, 1:400) and Donkey anti-Goat AF555 (Life Technologies A21432, 1:400).

Samples were washed in Ce3D washing buffer for 1 day. Samples were incubated in Ce3D clearing solution (N-methylacetamide/Histodenz/Thioglycerol) for 1 hour before mounting in fresh clearing solution. All preparations were scanned using a Leica SP8 WLL Upright scanning confocal including 405nm and White Light Laser (470-670nm) for excitation and imaging with 20X Apo long working distance water immersion objectives. Z steps were acquired every 5 mm. Ce3D protocol: PMID 28808033

For image analysis and quantification, z-stack images were rendered in 3D dimensions and quantitatively analyzed using Bitplane Imaris v9.8 software package (Andor Technology PLC, Belfast, N. Ireland). Individual CD8+ lymphocytes were annotated using the Imaris surfaces function based on the fluorescent signal (based on antibody staining), along with additional co-stains (e.g., CD45 IV, RFP) and size/morphological characteristics. 3D reconstructions of alpha smooth muscle actin (aSMA)-labeled structures and Iba+ cells were performed using Imaris surface function. Three dimensional distances between CD8+ lymphocyte subsets and Iba1+ surfaces were calculated using the Imaris object-object function.

### Cell Culture

To test the functional consequences of fibroblast deletion of *Tnfaip3* in vitro, freshly isolated lung fibroblasts from *Tnfaip3^flox/flox^* mice were cultured in DMEM/F-12 (Thermo Fisher, 11330032) with 10% FBS and 1% Pen/Strep. The medium was changed every 2 days and lung fibroblasts were maintained for no more than 3 passages. The fibroblasts were infected with Adenovirus-cre recombinase (UI Viral Vector Core, Ad5CMVCre #VVC-U of Iowa-5) or Adenovirus-empty (UI Viral Vector Core, Ad5CMVEmpty, #VVC-U of Iowa-272) at a titer of 0.5 × 10^8 pfu/100,000 fibroblasts. The following day cells were washed twice with PBS and fresh media was added to the wells. 3 days after viral infection cells (or media) were either utilized for further in vitro assays or lysed with Qiagen RLT buffer (Qiagen, 74104) and stored at -80°C in preparation for downstream gene expression analysis.

For fibroblast-T cell co-cultures, wildtype B6 splenic T cells were isolated by negative selection with the EasySep Mouse T cell Isolation Kit (Stemcell Technologies, #19851) according to manufacturer protocol, and 1 × 10^5 T cells were then added to co-culture with 2 × 10^4 Adenovirus-cre recombinase or Adenovirus-empty treated *Tnfaip3^flox/flox^* fibroblasts in T cell media (Advanced RPMI 1640 (Gibco, 12633012) with 10%FBS, 1%P/S, 1x GlutaMAX (Gibco, 35050061), 50 µM β-mercaptoethanol and 10ng/mL recombinant mouse IL-7 (R&D, 407-ML-005/CF)) in a 24-well plate. After 4 days of co-culture cells were lifted with TrypLE (Gibco, 12563011), and stained with anti-CD3-FITC and anti-PD-1-PE-Cy7 antibodies as described under “Tissue Dissociation and Flow Cytometry” prior to analysis. CountBright Absolute Counting Beads (Invitrogen, C36950) were used to determine absolute cell numbers.

To assess the ability of various cytokines to promote the formation of *Gzmk*+ CD8+ T cells, either resident (ivCD45-CD45+CD11b-CD3+) or circulating (ivCD45+CD45+CD11b-CD3+) cells were isolated from GATOR mice and 2.75 × 10^4 cells were plated in a 96-well round bottom plate in T cell media +/- supplementation with recombinant mouse IL-12 (15 ng/mL, R&D, 419-ML-010/CF) or recombinant mouse IL-15 (100 ng/mL, R&D, 447-ML-010/CF). After 5 days of culture cells were stained with anti-CD3-AF488, anti-CD4-BV711, anti-CD8-BV786 and DRAQ7 (live/dead) and analyzed by flow cytometry. Endogenous tdTomato expression from the GATOR mouse was used to assess *Gzmk* expression within T cell subsets. CountBright Absolute Counting Beads (Invitrogen, C36950) were used to determine absolute cell numbers.

For macrophage-T cell co-cultures, interstitial macrophages (ivCD45-CD45+Ly6g-SiglecF-CD11b+) were sorted from wildtype B6 mice and plated at a concentration of 5 × 10^4 cells/96-well flat bottom TC-treated plate in T cell media with 10ng/mL recombinant mouse MCSF (R&D, 416-ML-010/CF). After 2 days of culture any non-adherent cells were removed and 1×10^5 splenic T cells from GATOR mice, isolated by negative selection (StemCell Technologies, 19851), were added to each well +/-rat IgG2a (BioXCell, BE0089-R001mg) or rat anti-mouse IL-15RB (BioXCell, BE0298-R001mg) at a concentration of 20 ug/mL. After 5 days of co-culture, cells were lifted with TrypLE (Gibco, 12563011) and stained with anti-CD3-AF488, anti-CD4-BV711, anti-CD8-BV786 and DRAQ7 and analyzed by flow cytometry. Endogenous tdTomato expression from the GATOR mouse was used to assess *Gzmk* expression within T cell subsets. CountBright Absolute Counting Beads (Invitrogen, C36950) were used to determine absolute cell numbers.

In vitro stimulation of sorted lung resident GZMK+ (ivCD45-CD45+CD11b-CD3+CD8+tdTomato+) or GZMK-(ivCD45-CD45+CD11b-CD3+CD8+tdTomato-) CD8+ T cells was performed by plating between 5500 and 9000 cells (depending on experiment) per well of a 96-well round-bottom plate in 200uL T cell media with 1uL of anti-CD3/anti-CD28 mouse T cell activating beads (Thermo Fisher, 11456D). After 4 days of culture cells were stained with anti-CD3-FITC (Invitrogen, 11003282, clone 17A2), anti-CD8-BV786 (BD Biosciences, 563332, clone 54-6.7) and DRAQ7 viability dye and analyzed by flow cytometry using CountBright Absolute Counting Beads to assess cell numbers. In studies where intracellular IFNg levels was assessed, Golgiplug (BD Biosciences, 555029) was added to the media 3 hours prior to cell collection and cells were fixed and permeabilized with the Foxp3 Staining Buffer Set (eBioscience) prior to intracellular staining with anti-IFNg-APC (Invitrogen, 17-7311-81, clone XMG1.2). Supernatant was collected and stored at -80°C for downstream protein analysis.

For in vitro culture of lung fibroblasts or bone marrow derived macrophages (BMDMs), cells were prepared as follows. For fibroblasts, lung cells were harvested and lineage (CD45+, CD31+, EpCAM+) cells were removed with magnetic bead depletion as described under “Tissue Dissociation and Flow Cytometry”. Cells from one mouse were plated in a 10 cm TC-treated dish in fibroblast media (DMEM/F-12 with 10% FBS and 1%Pen/Strep). After 2-3 days of culture, adherent fibroblasts were lifted with TrypLE (Gibco, 12563011) and replated at a density of 4 × 10^4 cells per 48 well TC-treated dish. To grow bone marrow derived macrophages (BMDMs), following euthanasia mouse femurs and tibias were removed and crushed with a mortar and pestle to release the bone marrow. Cells were passed through a 70 μm filter and centrifuged at 350g x 5 minutes prior to RBC lysis x 2 minutes, followed by dilution with FACS buffer and 40 μm filtration prior to repeat centrifugation and cell counting. 10 × 10^6 bone marrow cells were plated per 10 cm non-TC treated dish in macrophage media (Advanced RPMI1640 with 10% FBS, 1x GlutaMAX, 1x P/S, 50 µM β-mercaptoethanol and recombinant mouse MCSF at 20ng/mL (R&D, 416-ML-010/CF)). Cells were fed on day 4 of growth with additional media. On day 7 non-adherent cells were removed, and adherent macrophages were lifted with cold PBS +5 mM EDTA and replated in macrophage media at a density of 6 × 10^4 cells per 48 well TC-treated dish. The day after replating, cell media from fibroblasts and BMDMs was replated with fresh media +/-recombinant mouse GZMK (MyBioSource, MBS2029386) at 100ng/mL. Per company communications, endotoxin levels of GZMK were measured by the LAL method and found to be <1.0EU per 1ug. After 3 hours of stimulation, cells were washed with PBS, lysed in Qiagen RLT buffer (QIAGEN, 74106), and stored at -80°C until downstream analysis.

### ELISA quantification of IFNg

Supernatant from T cells stimulated in vitro with anti-CD3/CD28 activating beads (see above under “Cell Culture”) was analyzed for IFNg levels with the Mouse IFN-gamma Quantikine ELISA Kit (R&D, MIF00-1) according to manufacturer protocol. Samples were read on a BioTek Synergy H1 Plate Reader.

### Luminex cytokine assay

For protein concentration from lung homogenates, samples were analyzed by Eve Technologies (https://www.evetechnologies.com/) using their 32-Plex Discovery Assay on the Luminex 200 platform. To prepare samples for analysis the right lower lobe was flash frozen on dry ice at time of collection and stored at -80°C. Tissues were then thawed on ice, and the weight of each lobe was recorded. Samples were then transferred to a bead homogenizer tube (Benchmark #D1032-30: 2 mL tubes, 3.0mm high impact zirconium beads) and 1x RIPA buffer (CST #980) containing 1mM PMSF (Cell Signaling Technologies, #8553) was added to each tube at a volume of 10ul per mg of tissue. Tissues were then homogenized using the Precellys Evolution tough Homogenizer with the following settings (2×30sec, 30sec pause, 6500 RPM, 4°C). Dry ice was added to the top compartment to keep the samples cold during processing.

Lysate was spun at 10,000g × 10 minutes at 4°C and supernatants were subsequently transferred to a new tube. Protein concentration of supernatant was measured using the Pierce BCA Assay Kit (Thermo Fisher, 23227). Samples were diluted to a final concentration of 2mg/mL which were stored at -80°C and then shipped to Eve Techologies on dry ice for analysis. Using initial tissue weights, reported protein concentrations from Luminex assay were back-calculated to determine original tissue concentrations and reported in the manuscript as picogram of protein/milligram of tissue.

### Transwell Migration Assay

For T cell and monocyte transwell migration assays, media from Adenovirus-cre recombinase vs adenovirus-empty treated fibroblasts was conditioned for 48 hours and then transferred to the bottom well of a 24-well transwell plate with 5um pores (Fisher Scientific 07-200-149). In the upper well either 3 × 10^5 wildtype B6 splenic T cells or 2 × 10^5 wildtype B6 bone marrow monocytes were added in unconditioned T cell media. After incubation of T cells or monocytes for 3 hours, cells that had migrated to the bottom chamber were collected and numbers of CD3+ (T cells) or CD11b+ (monocytes) cells were analyzed by flow cytometry along with DAPI as a viability dye. CountBright Absolute Counting Beads (Invitrogen, C36950) were used to determine absolute cell numbers.

### Quantitative RT-PCR (qPCR)

Total RNA was obtained from cells using either the PicoPure RNA Isolation Kit (Applied Biosystems, KIT0204) for sorted cells, or the RNeasy mini kit (QIAGEN, 74106) for cultured cells following the manufacturers’ protocols. cDNA was synthesized from total RNA using the SuperScript Strand Synthesis System (Thermo Fisher, 18080044). Quantitative RT-PCR (qRT-PCR) was performed using the SYBR Green system (Thermo Fisher, F415L). Relative gene expression levels after qRT-PCR were defined using the ΔΔCt method and normalizing to the housekeeping genes. The qRT-PCR primers used for mouse are as follows: Ccl2-F: ACCTGCTGCTACTCATTCACC, Ccl2-R: ATTCCTTCTTGGGGTCAGCA, Ccl7-F: CCATGAGGATCTCTGCCACG, Ccl7-R: GCAGCATGTGGATGCATTGG, Ccl8-F: TCTACGCAGTGCTTCTTTGCC, Ccl8-R: AAGGGGGATCTTCAGCTTTAGTA, Csf1-F: GTGTCAGAACACTGTAGCCAC, Csf1-R: TCAAAGGCAATCTGGCATGAAG, Csf3-F: CCCACCTTGGACTTGCTTCA, Csf3-R: GGAAGGCAGAAGTGAAGGCT, Cxcl1-F: ACTCAAGAATGGTCGCGAGG, Cxcl1-R: GTGCCATCAGAGCAGTCTGT, Cxcl2-F: CACTCTCAAGGGCGGTCAAA, Cxcl2-R: TGGTTCTTCCGTTGAGGGAC, Gapdh-F: GGCCCCTCCTGTTATTATGGGGGT; Gapdh-R: CCCCAGCAAGGACACTGAGCAAGA; Gzmk-F: TGGCTGGCGTTTATATGTCTTC Gzmk-R: GCTGCGGTACTGGATGGAC, Il1b-F: TGCCACCTTTTGACAGTGATG, Il1b-R: TGATGTGCTGCTGCGAGATT, Il6-F: AACGATGATGCACTTGCAGA, Il6-R: CTCTGAAGGACTCTGGCTTTG, p16INK4a-F: AATCTCCGCGAGGAAAGC; p16INK4a-R: GTCTGCAGCGGACTCCAT; Tnf-F: CTGAACTTCGGGGTGATCGG, Tnf-R: GGCTTGTCACTCGAATTTTGAGA, Tnfaip3-F: TGTCTCCTTAAGGGTGCTGC, Tnfaip3-R: TCGTGAAGTCAGGAAGCTCG.

### Quantification and statistical analysis

All data points represent measurements of an individual mouse, or for in vitro experiments, an individual well. GraphPad Prism was used for all statistical analyses. Statistical significance was determined by ordinary one-way ANOVA, two-way ANOVA or one or two-tailed Student’s t-test as indicated in figure legends. Error bars represent standard deviation. For consistency in these comparisons, the following denotes significance in all figures: \**P* < 0.05, \*\**P* < 0.01, \*\*\**P* < 0.001, \*\*\*\**P* < 0.0001.

## Data availability

The GEO datasets presented in the manuscript are GSE286085 (Xenium), GSE285880 (Bulk fibroblast and Resident Immune scRNAseq), GSE286324 (CD3 TCR scRNAseq) and GSE286988 (Aged vs young lung fibroblast scRNAseq).

## Code availability

No custom codes were developed and used in this manuscript. All codes are available by request to the corresponding author.

